# An Asgard archaeon with internal membrane compartments

**DOI:** 10.1101/2025.11.06.686947

**Authors:** Fraser I. MacLeod, Andriko von Kügelgen, Magdalena K. Lechowska, Joe Parham, Iain A. Richard, Emily J. Aguilar-Pine, Brendan P. Burns, Brett J. Baker, Tanmay A.M. Bharat, Buzz Baum

## Abstract

The emergence of eukaryotes from a merger between an archaeon and a bacterial cell ∼two billion years ago involved a profound change in cellular organisation. While the order in which different features of the eukaryotic cell arose remains a matter of controversy^1-3^, close archaeal relatives of eukaryotes have recently been identified that possess homologues of eukaryotic trafficking machinery^4,5^ and a complex cell architecture^6,7^. The members of this phylum, the Asgard archaea (*syn*. Prometheoarchaeota) described so far, however, lack internal membrane-bound compartments, and therefore have shed little light on origins of the hallmark eukaryotic endomembrane system. Here we report the cell biological analysis of a member of the Heimdallarchaeia, *Candidatus ‘*Y. umbracryptum’ in enriched mixed microbial communities. Possessing a small genome encoding few homologues of eukaryotic membrane remodelling machinery^8^, *Ca.* Y. umbracryptum cells in late-stage cultures resemble previously described Asgard archaea with extensive cellular protrusions. Surprisingly, during early stages of culture growth *Ca.* Y. umbracryptum cells have fewer protrusions but possess numerous intracellular vesicles, most of which have a luminal surface that morphologically resembles the outer coat of the plasma membrane. These data alter our view of eukaryogenesis by identifying a close archaeal relative of eukaryotes with an endomembrane system, whose appearance changes with cell state.

## Introduction

The emergence of eukaryotes about two billion years ago was associated with a profound change in cellular organisation as comparatively, ultrastructurally simple archaea and bacteria gave rise to complex eukaryotic cells that possess numerous different internal membrane-bound compartments^2^. These include the nucleus, mitochondria, the endoplasmic reticulum, the Golgi apparatus, lysosomes, along with intracellular vesicles that mediate the transport of lipids and protein cargo between these different organelles. In addition, subsets of these internal vesicles traffic to and from the plasma membrane^9^, mediating communication between the cell interior and the external world.

The origins of this complex architecture of the eukaryotic cell and the order in which the different compartments arose remain unknown^2^. Nevertheless, as the result of phylogenetic analyses^10,11^ and structural data^12^, it is now widely accepted that eukaryotes emerged from a process of endosymbiosis when a cell related to modern Alphaproteobacteria took up residence inside a host cell to give rise to mitochondria. While the identity of the host cell long remained a point of contention, recent phylogenetic studies using a range of marker genes of archaeal origin firmly place eukaryotes within the Asgardarchaeota (*syn*. Prometheoarchaeota). More specifically, most studies place eukaryotes within^4,8,13^ or as sisters to^14^ the Heimdallarchaeia. In support of this conclusion, the genomes of many Asgard archaea also contain genes with homologues in eukaryotes that function in membrane remodelling^8,15^. However, with the possible exception of ESCRT-III polymers^4,16-19^ and the actin machinery from Asgard archaea^20,21^, which exhibit structurally and functionally similar properties to their eukaryotic counterparts when studied *in vitro*, it is unclear whether these perform analogous roles in archaea. Similarly, while the first images of Asgard lineage cells revealed a number of cellular hallmarks of eukaryotes, including protrusions rich in an actin cytoskeleton^7^, the Asgard archaea imaged thus far lack clear evidence of internal membrane-bound compartments. As a result, it has not yet been possible to distinguish between models of the evolution of eukaryotes that differ in their predictions about the relative timing of the emergence of different parts of the endomembrane system^1,3,22,23^.

In this article, we report the cell biological analysis of a member of the Heimdallarchaeia, *Candidatus* ‘Y. umbracryptum’, which was cultivated as part of a mixed microbial community from a sample taken from microbial mats in Shark Bay, Western Australia. Strikingly, *Ca.* Y. umbracryptum cells possess a diverse set of intracellular vesicles, most of which appear empty at the resolution of cryogenic electron tomography (cryo-ET), which are lined with a surface coat that is morphologically and ultrastructurally indistinguishable from the coat on the external face of the plasma membrane. Furthermore, *Ca.* Y. umbracryptum cells were observed wrapping extracellular material, and in one case another organism, in the mixed culture. Since vesicles were only present in high numbers in cells taken from early-stage cultures that possess markedly fewer protrusions, their formation and loss appears to be something that cells regulate depending on environmental circumstances. Taken together, these data suggest the possibility that *Ca.* Y. umbracryptum has a dynamic endomembrane system and can likely both internalise material from the environment, a prerequisite for the final steps in the process leading to the acquisition of mitochondria, and traffic intracellular material to the cell surface. Thus, by reporting the cell biology of a member of the Heimdallarchaeia, an archaeon with an abundance of internal vesicles, this study further narrows the divide that separates archaea and eukaryotes^4^ and sheds new light on the emergence of the endomembrane system, which is a hallmark of eukaryotes.

## Results

### Heimdallarchaeia enrichment cultures

Microbial mat samples (**Fig. 1a**) rich in Asgard archaea (between approximately 2.5% and 5% of DNA sequencing reads) were collected from the Nilemah tidal flats of Hamelin Pool in Shark Bay, Western Australia as described previously^24,25^, and used to inoculate anaerobic cultures (Materials and Methods). By passaging these cultures over six years in different growth media, mixed cultures were identified that, based on 16S rRNA gene sequences, contained a single Heimdallarchaeia lineage at a stable level of 25-30% of the population, along with a set of eight other archaea and bacteria, including uncharacterised species of Bacteroidia, Desulfovibrionia, Halanaerobiia, Bathyarchaeia, and Thermoplasmata (**Fig. 1b**). We named the Heimdallarchaeia lineage Asgard archaeon, *Candidatus* ‘Y. umbracryptum’. Based upon 16S rRNA qPCR, we estimate that *Ca.* Y. umbracryptum has a minimum doubling time of about three days and an average doubling time of seven and a half days during log phase growth (**Fig. 1c**). Short-read DNA sequencing of these cultures enabled the assembly of a near-complete Heimdallarchaeia genome in 55 contigs (**Supplementary Table 1**). A maximum-likelihood phylogenetic analysis using recently published marker sets^8,26^ identified

**Figure 1:**
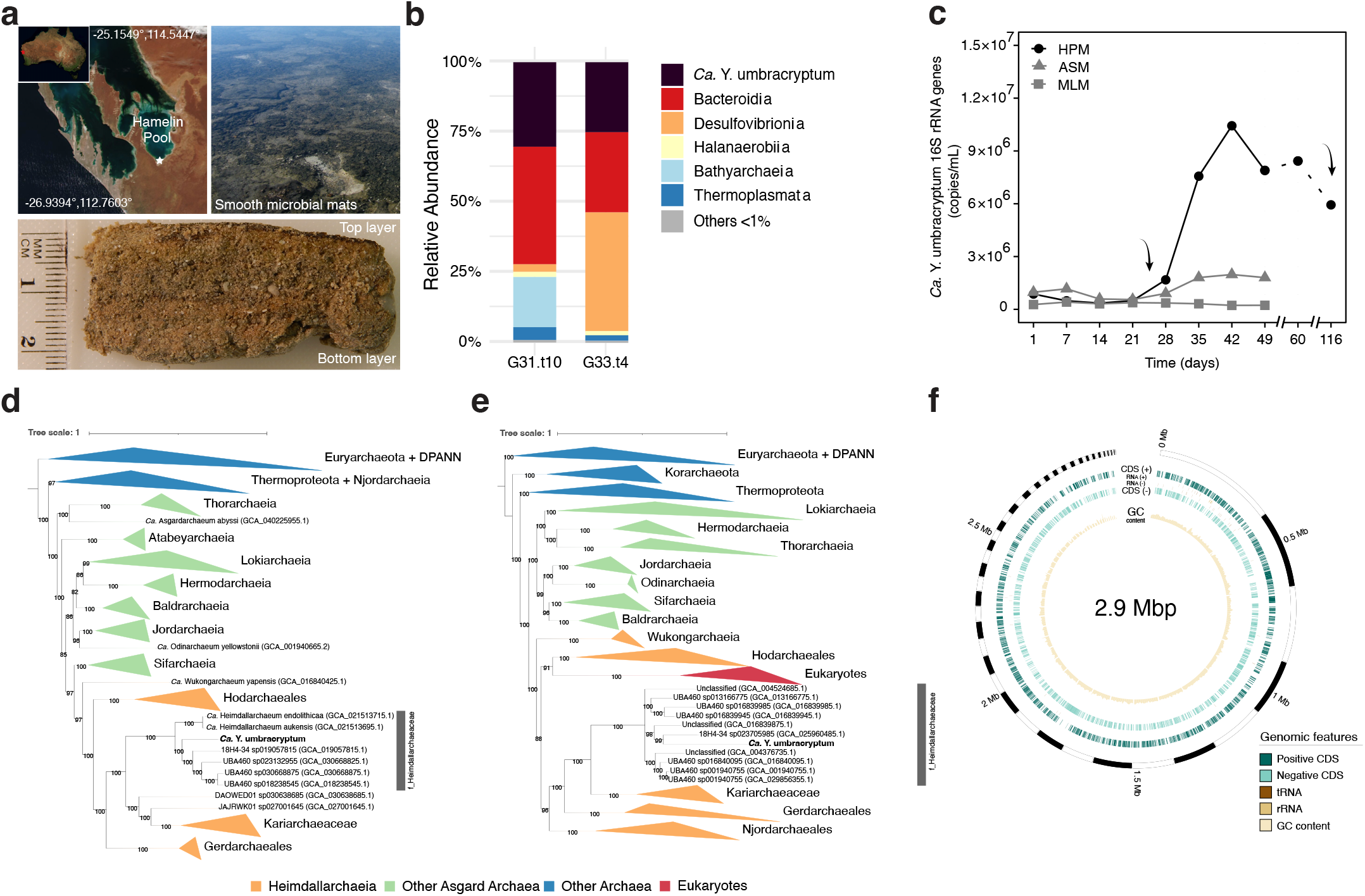
Isolation and cultivation of Heimdallarchaeaceae *Candidatus* Y. umbracryptum. **(a)** Satellite image of Australia showing the location of Shark Bay and satellite image of Shark Bay showing the location of the sample site, taken on day of sampling (top left). Satellite images were retrieved from NASA’s Earth Science Data and Information System (ESDIS). Image of the smooth microbial mats in the Hamelin Pool inter-tidal zone (top left). Image of the smooth microbial mat sample used as the initial inoculum (bottom). **(b)** Relative abundance of *Ca*. Y. umbracryptum in late stage (G31.t10) and early stage (G33.t4) enrichment cultures, as determined by amplicon sequencing using the universal primer set 926F-1392wR. **(c)** Growth of *Ca*. Y. umbracryptum as determined by qPCR under different media conditions. Hamelin Pool Medium (HPM), Archaeal Salt Medium (ASM), Minimal Loki Medium (MLM)^7^. Early and late sampling timepoints for light-mi-croscopy and cryo-ET experiments are indicated by curved black arrows. **(d)** Maximum-likelihood phylogenetic placement of *Ca*. Y. umbracryptum within the archaea using ar53 marker genes from GTDB-tk^26^. **(e)** Maximum-likelihood phylogenetic placement of *Ca*. Y. umbracryptum including the placement of eukaryotes using NM57 marker genes^8^. **(f)** Schematic of the *Ca*. Y. umbracryptum draft genome, illustrating genome size, GC content, the set of tRNA and rRNA included in the genome, and the binning of the genome into 55 contigs.

this organism as a member of the class Heimdallarchaeia and family Heimdallarchaeaceae (**Fig. 1d, Supplementary Fig. 1**) and a close relative of eukaryotes (**Fig. 1e and Supplementary Fig. 1**).

The genome of *Ca.* Y. umbracryptum, while typical for a member of the family Heimdallarchaeaceae, was markedly smaller than those reported for the recently cultivated Lokiarchaeia species *Prometheoarchaeum syntrophicum*^6^, *Candidatus* ‘Lokiarchaeum ossiferum‘^7^ and Hodarchaeales^27^ (**Fig. 1f and Supplementary Fig. 2a**). In line with this, its genome encodes fewer proteins associated with a host of functions from information processing to metabolism, cellular ultrastructure and division than the other cultivated Asgard archaea (**Supplementary Fig. 2b**). Most strikingly, the reconstruction of its metabolic pathways (**Supplementary Fig. 3a-b**) revealed that *Ca*. Y. umbracryptum is only capable of synthesizing six amino acids (alanine, methionine, glutamic acid, serine, glutamine, glycine). Since it also lacks degradation pathways for most amino acids, these data suggest the possibility that raw material for protein synthesis in *Ca.* Y. umbracryptum is supplied from external sources of amino acids or peptides. The *Ca.* Y. umbracryptum genome also encodes multiple ATP-binding cassette transporters that includes a system for small peptide uptake. In other respects, however, its complement of genes seemed largely complete. The genome assembly encodes 45 tRNAs (**Fig. 1f and Supplementary Table 1**), along with the genes required to perform tRNA aminoacylation for most amino acids (**Supplementary Fig. 3a**), as well as the genes required to perform basic nucleic acid interconversions, and key steps of nucleic acid biosynthesis/degradation. A nearly complete archaeal-type mevalonate (MVA) pathway was also present (**Supplementary Fig. 3a**), suggesting that *Ca.* Y. umbracryptum can synthesize necessary precursors for any isoprenoid-based compounds, including archaeal membrane lipids^28^. These data support the idea that *Ca.* Y. umbracryptum depends on partners or extracellular material for growth as was suggested to be the case for other Asgard archaea in mixed cultures^6,7.^ We also noted that while the *Ca.* Y. umbracryptum genome encodes homologues of eukaryotic actin and ESCRT-I,-II, and-III proteins (**Supplementary Fig. 3b and 3c**) and 14 Arf/SAR type GTPase domain proteins^4^, it lacks clear homologues of proposed membrane remodelling systems reported in other Asgard genomes^8,13^, such as Sec23/24, TRAPP, SNAREs and arrestins (**Supplementary Fig. 3c**).

The paucity of “eukaryotic signature proteins” (ESPs) present in its genome relative to the genomes of Lokiarchaeia species *P. syntrophicum*^6^, *Ca.* ‘Lokiarchaeum ossiferum‘^7^ and Hodarchaeales^27^ led us to examine the cell biology of *Ca.* Y. umbracryptum. To study its morphology using light microscopy, cells from late-stage cultures were visualised under anaerobic conditions on a confocal microscope. Under these conditions, alongside filamentous bacteria and rare cells with distinct morphologies (**Supplementary Fig. 4**), we observed cells similar in appearance to the Asgard archaea described in recent reports^6,7,25,27,29,30^, with a central cell body containing the DNA and long branched protrusions (**Fig. 2a**).

**Figure 2:**
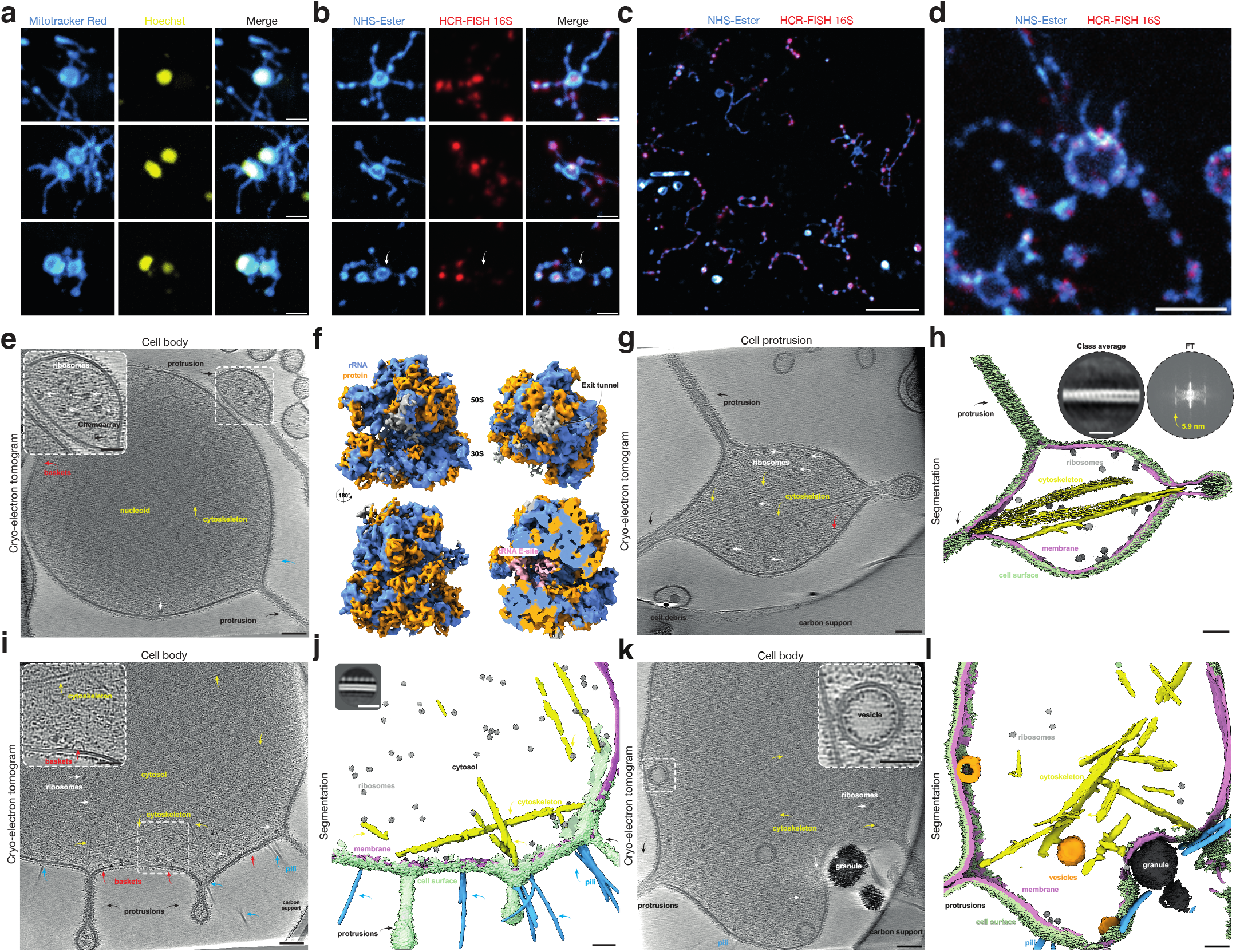
Light microscopy and cryo-ET of stationary phase *Ca.* Y. umbracryptum cultures. **(a)** Examples of cells from live-cell imaging of the stationary culture. MitoTracker Red CMXRos (depicted in cyan hot) was used as a cell contour stain and Hoechst 33342 (yellow) as a DNA stain in two biological replicates. Scale bars: 1 µm. **(b)** Example images of cells from cultures labelled with *Ca.* Y. umbracryptum-specific HCR-FISH 16S rRNA probes (red) and with NHS-ester (cyan hot) as a generic cell contour stain from two biological replicates. White arrow depicts cell with HCR-FISH signal in protrusions but not in the cell body. Scale bars: 1 µm. **(c)** Overview image of HCR-FISH data shown in b), Scale bar: 5 µm. **(d)** Expansion HCR-FISH image of *Ca.* Y. umbracryptum with HCR-FISH 16S probes (red) with NHS-ester (cyan hot) from two biological replicates. Scale bar: 5 µm (see also Supplementary Fig. 4). **(e)** Cryo-ET slice through the cell body of (putative) *Ca.* Y. umbracryptum with few ribosomes (white arrows) and cytoskeletal filaments (yellow arrows). Inset shows close up of cellular protrusion filled with multiple ribosomes (white). Scale bar: 100 nm; Inset scale bar: 50 nm (see also Supplementary Fig. 5). **(f)** Subtomogram averaging structure of the ribosome from these cells at ∼10 Å (see also Supplementary Fig. 6a-c). **(g)** Cryo-ET slice through a cell protrusion showing multiple cytoskeletal filaments (yellow arrows). Scale bar: 100 nm. **(h)** Semantic segmentation of the tomogram shown in panel g) depicting the cell membrane (pink), cell-surface (pale green), ribosomes (grey) and cytoskeletal filaments (yellow). Inset shows two-dimensional class average of cytoskeletal filaments and the corresponding Fourier Transform (FT, power spectrum shown) with a repeat at ∼5.9 nm. Scale bar: 100 nm; Inset scale bar: 25 nm (see also Supplementary Fig. 6d). **(i)** Cryo-ET slice through the cell body of *Ca.* Y. umbracryptum with multiple cytoskeletal filaments (yellow). Inset shows close up of cytoskeletal filaments and a membrane attached basket. Scale bar: 100 nm; Inset scale bar: 50 nm. **(j)** Semantic segmentation of the tomogram shown in panel i) with similar color schemes as panel h), but including pili (blue). Inset shows a two-dimensional class average of the cell membrane. Scale bar: 100 nm. Inset scale bar: 25 nm (see also Supplementary Video 1). **(k)** Cryo-ET slice through a cell body of a *Ca.* Y. umbracryptum cell with multiple intracellular vesicles. Inset shows a close-up of one intracellular vesicle with clear separation of the two leaflets of the lipid bilayer. Scale bar: 100 nm; Inset scale bar: 50 nm. (see also Supplementary Fig. 7 and Supplementary Video 2). **(l)** Semantic segmentation of the tomogram shown in panel k) with similar colour schemes as panel h) but including versicles (orange) and extracellular granules (black). Tomograms: *n*=135, from two biological replicates.

To confirm the identity of *Ca.* Y. umbracryptum in these mixed cultures, we modified the imaging protocol for hybridised chain reaction fluorescent *in situ* hybridisation (HCR-FISH). Fixed and permeabilised cells were labelled with HCR-FISH probes specific to the *Ca.* Y. umbracryptum 16S rRNA gene and a fluorescent NHS-ester to mark the contour of all the cells in the community (**Figs. 2b-c**). As expected, the *Ca.* Y. umbracryptum-specific 16S rRNA probe specifically labelled cells with long protrusions, many of which appeared to be part of extended networks, but did not label cells in Heimdallarchaeia-free cultures (**Supplementary Fig. 4b**). Conversely, the Heimdall cultures were not labelled by an HCR-FISH probe designed to complement Lokiarchaeia rRNA. Having confirmed the identity of these cells as *Ca.* Y. umbracryptum (**Fig. 2b and Supplementary Fig. 4**), to image the localisation pattern of rRNA at higher resolution, we processed the *Ca.* Y. umbracryptum-containing multicellular cultures for expansion microscopy by adapting a previously described protocol^31^ that uses a crosslinking agent to enhance the RNA retention, and observed cells after a 4X expansion using the same Heimdall-specific HCR-FISH probes. Again, this revealed networks of cells with elaborate protrusions rich in rRNA (**Fig. 2d**), similar those seen in other Asgard archaeal cultures^7,29^.

### Ultrastructural analysis of *Ca.* Y. umbracryptum

Having used HCR-FISH to confirm the identity of *Ca.* Y. umbracryptum cells in mixed communities, cryo-ET was employed to examine their ultrastructure. For this analysis, cells from late-stage cultures were collected from anaerobic serum bottles, applied directly onto electron microscopy grids and vitrified. In line with data from the light microscopy experiments, cells on the grids tended to have a small cell body with long undulating cell-surface protrusions (**Fig. 2e and Supplementary Fig. 5**). In tomograms, ribosomes could be seen that were abundant in protrusions, as was previously observed in *Ca.* L. ossiferum cultures^7^ but tended to be excluded from cell bodies that contained a dense nucleoid (**Fig. 2e**), in agreement with our HCR-FISH data (**Fig. 2b-d**). Using subtomogram averaging^32,33^ we generated a 10 Å-resolution structure of the ribosome in these cells (**Fig. 2f, Supplementary Fig. 6 and Supplementary Table 2**) that could be compared with previously reported atomic structures of an archaeal ribosome from *Thermococcus kodakerensis*^34^ and a bacterial ribosome^35^. The comparison confirmed the identity of these cells as archaea (**Supplementary Fig. 6**), in agreement with the HCR-FISH data. These cells also contained numerous linear cytoskeletal elements, many of which were bundled (**Fig. 2g**), especially within the thin necks connecting protrusions (**Fig. 2g and Supplementary Fig. 5**). These resembled the bacterial and archaeal actin filaments that have been observed by cryo-ET previously^36,37.^ and subtomogram averaging revealed a double-protofilament architecture with a 5.9 nm repeat close to that of prokaryotic actin^37^ (**Figs. 2g-h**). At the same time, these filaments had only a weakly discernible long-range protofilament twist that is characteristic eukaryotic actin^38^. Since *Ca.* Y. umbracryptum is the only species in the mixed community containing a close actin homologue in its genome, these data, when taken together with the ribosome subtomogram averaging, and with their characteristic morphology observed in HCR FISH, confirmed the identity of these cells. By cryo-ET we were also able to observe the bounding membrane of cells and membrane associated structures in detail. Like the surface of *P. syntrophicum* and *Ca.* L. ossiferum^6,7^ cells, the surface of *Ca.* Y. umbracryptum cells appeared coated with an amorphous proteinaceous layer rather than a continuous, paracrystalline S-layer lattice that is characteristic of many archaea^39-41^. The plasma membrane was also decorated with occasional pili of one type (**Fig. 2i-j**). Additionally, in many tomograms we observed basket-like structures attached to the internal face of the *Ca.* Y. umbracryptum plasma membrane (**Figs. 2e,g,i and Supplementary Fig. 7**), often in clusters (**Supplementary Fig. 7b**). Since similar structures are formed by SPFH family of proteins in bacteria and eukaryotes^42,43^, we searched the genome of *Ca.* Y. umbracryptum for SPFH homologues and identified two SPFH-family proteins that contain the extended alpha-helical domain required to form a basket (**Supplementary Fig. 7**).

### Intracellular vesicles in the cytosol are enriched in early-stage cultures

By cryo-ET we also identified several cells containing a small number of internal vesicles (**Figs. 2k-l, Supplementary Figs. 8a-h**). To ensure that this was not an experimental artefact of our cryo-ET vitrification in ambient conditions, we performed vitrification under anaerobic conditions with identical results (**Supplementary Fig. 8i-l**). Since these vesicles were rare in high-density, stationary, late-stage cultures, we then repeated the analysis, looking at cells taken from cultures at earlier stages in the microbial community growth cycle. For this analysis we examined cultures that were just entering log phase growth (3-4 weeks after inoculation) (**Fig. 1c**). While these contained ∼30% Heimdallarchaeia as measured by 16S amplicon sequencing, when imaged live or following fixation we observed few cells with protrusions (**Fig. 3a and Supplementary Fig. 4a**). Instead, these early-stage cultures contained a high proportion of cells that were spherical, slightly larger than the *Ca.* Y. umbracryptum cell bodies seen in late-stage cultures; a proportion of which had poorly resolved spot-like structures that labelled with the membrane probe (**Figs. 3a-b**). Using cryo-ET to examine the ultrastructure of these cells, we were able to confirm their identity as *Ca.* Y. umbracryptum based on the presence of actin-like filaments, their characteristic plasma membrane surface coat, and the ribosomal structure generated by subtomogram averaging that matched the structure from the late stage cells (**Fig. 3 and Supplementary Fig. 9**). Strikingly, most cells contained vesicles, often in very large numbers (**Figs. 3c-f and Supplementary Fig. 10**). While these vesicles had a range of morphologies, the major class seen across multiple biological replicates resembled those observed in late-stage cultures in shape, size and architecture (**Fig. 2k and Supplementary Fig. 8**). The majority were spherical and had diameters of between 60 and 90 nm (**Figs. 4a-b**). Most were positioned close to the cytoplasmic membrane without being topologically connected to it, and were devoid of dense contents. On their luminal side, the vesicular coat appeared morphologically indistinguishable from the cell surface (**Figs. 4a-d and 2j**). This characteristic coat ultrastructure was confirmed when we segmented and extracted all membranes (including the plasma membrane) observed in tomograms, and performed a sub-tomogram analysis by adapting previously described techniques^44^ (**Fig. 4d**). The presence of this internal coat was consistent with these vesicles being involved in material exchange with the surface membrane. In line with this hypothesis, in one instance a vesicle was identified that contained a pilus that was morphologically indistinguishable from pili observed on the cell surface (**Figs. 3d-f**) - implying that the vesicle was either in the process of delivering material to the surface or contained cell surface material that had been internalised.

**Figure 3:**
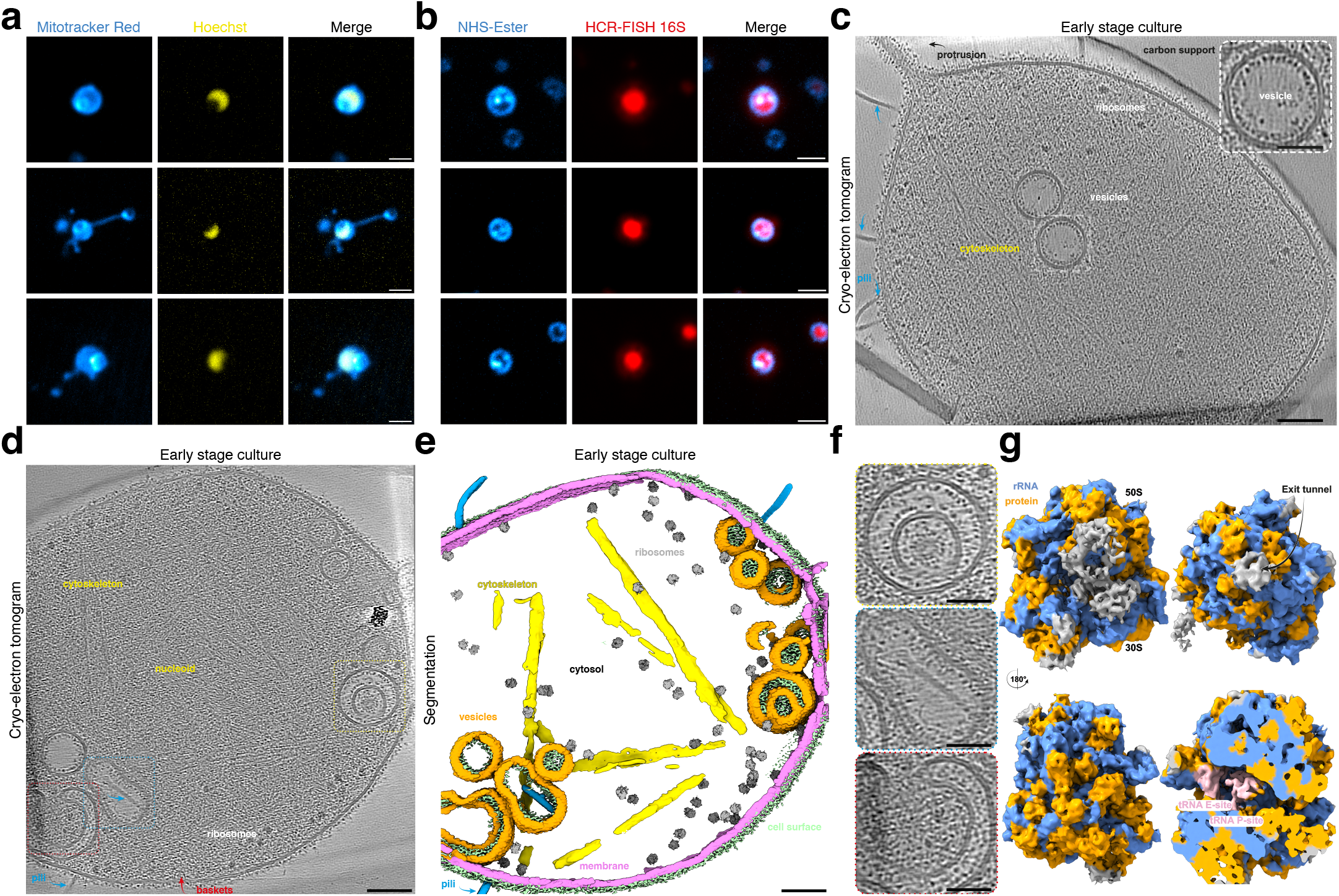
Light and electron microscopy of early phase *Ca.* Y. umbracryptum cultures. **(a)** Examples from live-cell imaging of the early log phase culture. MitoTracker Red CMXRos (depicted in cyan hot) was used to mark the *Ca.* Y. umbracryptum cell contour and Hoechst (yellow) the DNA in two biological replicates. MitoTracker Red CMXRos staining revealed large intracellular spheres of staining, which excluded Hoechst signal. Scale bars: 1 µm. **(b)** Example images of *Ca.* Y. umbracryptum cells labelled with specific HCR-FISH probes targeting the 16S rRNA (red) and NHS-ester (cyan hot) as a generic stain to mark the cell contour from two biological replicates (see also Supplementary Fig. 4). Cells identified as *Ca.* Y. umbracryptum by HCR-FISH labelling appeared to contain intracellular NHS-ester foci. **(c)** Cryo-ET slice through a *Ca.* Y. umbracryptum cell with few ribosomes, cytoskeletal filaments and intracellular vesicles marked. Inset shows close-up of one intracellular vesicle with clear separation of the two leaflets of the lipid bilayer. Scale bar: 100 nm; Inset scale bar: 50 nm. (see also Supplementary Figs. 9-10) **(d)** Cryo-ET slice through a cell body of early-log phase *Ca.* Y. umbracryptum with multiple intracellular vesicles. Scale bar: 100 nm. **(e)** Semantic segmentation of the tomogram shown in panel d) depicting the cell membrane (pink), cell-surface proteins (pale green), pili (blue), ribosomes (grey), cytoskeletal filaments (yellow) and intracellular vesicles (orange). Scale bar: 100 nm. (see also Supplementary Fig. 10 and Supplementary Video 3). **(f)** Close-up views of cryo-ET slices shown in panel d) depicting 1) closed multi-lamellar vesicle; 2) internalized pilus; and 3) open multi-lamellar vesicle. Scale bars: 50 nm. **(g)** Subtomogram averaging structure of the ribosome at ∼10.7 Å resolution. Tomograms: *n*=42, from two biological replicates.

Although the lumen of most vesicles was largely empty when viewed at this resolution, we also observed a population of intermediate-sized vesicles (75-150 nm) that contained smaller vesicles inside (**Figs. 4b,c,e**). In some cases, the inner membrane was connected to the outer vesicular membrane by three-way junctions (**Supplementary Fig. 10a,e**). In addition to this, we observed a number of larger vesicles which had a non-spherical shape. These included vesicles with donut-shaped form and, in one case, a double membrane that formed a C-shape (**Fig. 3d-f and Supplementary Fig. 10c**), reminiscent of an early stage intermediate of multivesicular body formation or autophagosome assembly in eukaryotes^45^. In this instance, it was clear that the inner compartment contained cytoplasm and ribosomes as would be expected if this process were catalysed by ESCRT-III dependent membrane remodelling^46^. We also observed several examples of extracellular material, including another archaeal cell, being wrapped within surface invaginations of the *Ca*. Y. umbracryptum cell (**Supplementary Fig. 11**).

**Figure 4:**
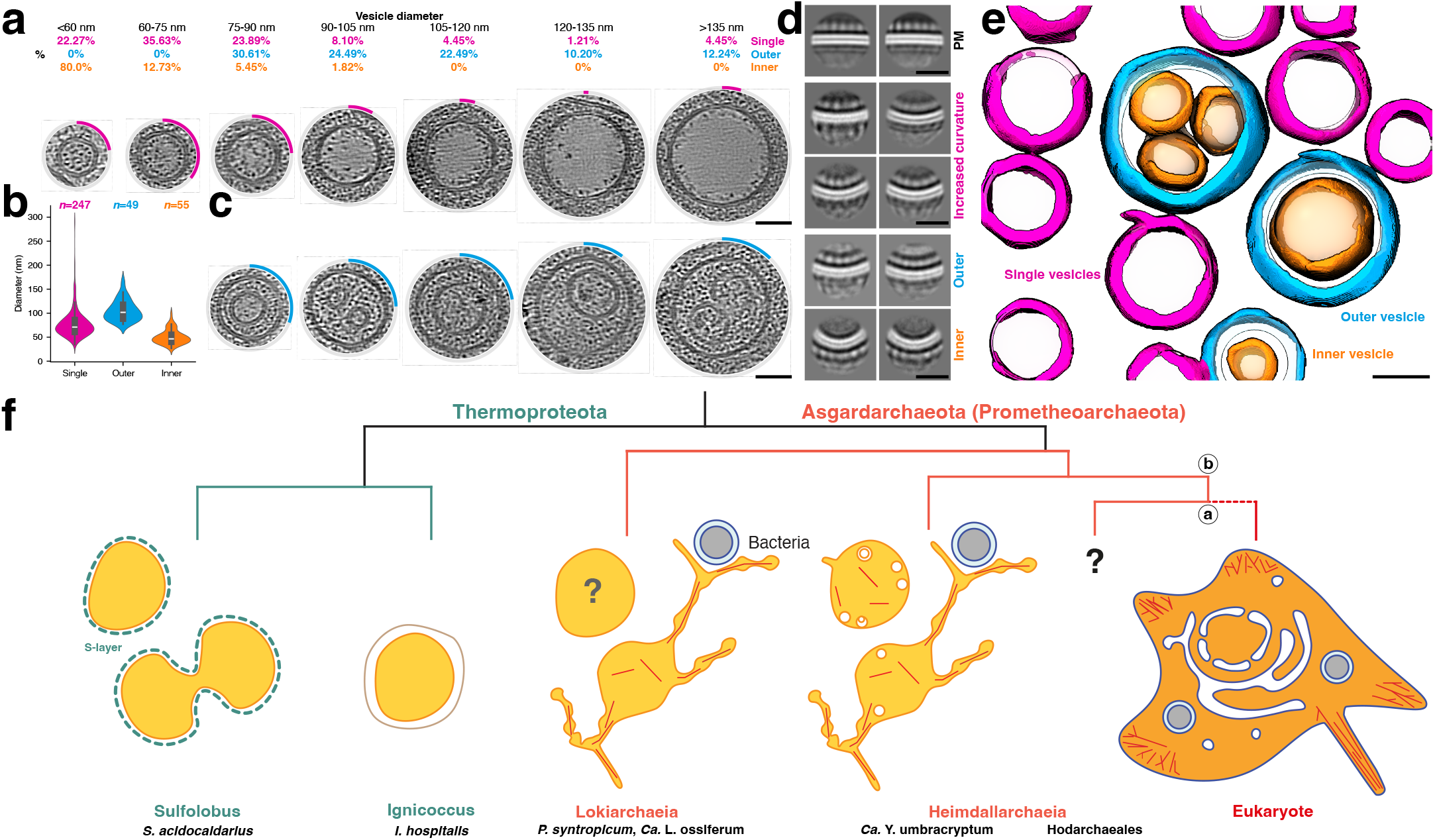
Intracellular vesicles of *Ca.* Y. umbracryptum. **(a)** Gallery of cryo-ET slices showing single vesicles arranged by increasing diameter from left to right. The percentages of intracellular vesicles are measured in groups from <60 nm to >135nm in bin-sizes of 15 nm. The percentage of each bin-size is additionally visualized as an arc (magenta) surrounding a representative vesicle of the corresponding diameter. Scale bar: 50 nm. **(b)** Violin plot of measured vesicle diameter (nm) of single (magenta), outer filled vesicles (cyan) and inner vesicles (orange). **(c)** Gallery of multi-lamellar vesicles ordered by diameter with the same group sizes and percentages displayed in panel a). Scale bar: 50 nm. **(d)** Subtomogram analysis of the plasma membrane (PM) and internalized membranes from vesicles produced two-dimensional averages at different membrane curvatures. The lipid bilayer (white) is clearly resolved in the averages, which is covered by an amorphous, proteinaceous surface coat in all cases. Two-dimensional averages of outer filled vesicles show an additional fuzzy lipid bilayer within its lumen, while averages of inner vesicle show inverted topology of the proteinaceous coat (i.e. coat on the outside, with positive curvature of the underlying membrane). Scale bars: 25 nm. **(e)** Semantic segmentation of a *Ca.* Y. umbracryptum cell with single (magenta), outer filled vesicles (cyan) and inner vesicles (orange), overlaid with a fitted transparent sphere in the same color. Scale bar: 50 nm. Tomograms: *n*=35, from two biological replicates. **(f)** Cartoon representation of cell morphology of Thermoproteota (light green), Asgardarchaeota (*syn.* Prometheoarchaeota - light red), diderm bacteria (pale grey) and Eukaryotes (red). Eukaryotes are depicted to branch within (a)^4,8,13^ or sister (b)^14^ to Heimdallarchaeia. The architecture of early-stage Lokiarchaeial cells remains unresolved (grey question mark). Cells are not depicted to scale.

Finally, we observed a cell with large internal compartments containing electron-dense material similar to that seen in the medium, which may have been internalised (**Supplementary Fig. 11f**).

## Discussion

This paper describes the cell biology of *Ca.* Y. umbracryptum, the first member of the Heimdallarchaeia to be cultivated and imaged. Like the other Asgard archaea that have been subjected to a detailed cell biological analysis^6,7^,29, when taken from dense, late-stage cultures, these cells were characterised by a central cell body that contains the genome from which long, branched protrusions extend. *Ca.* Y. umbracryptum (like other Asgard archaea) appears to lack many key enzymes in several core metabolic pathways consistent with suggestions^6^ that the protrusions facilitate syntropic interactions with partners in the mixed community in which they live, or assist in the absorption of metabolites from the environment. Unlike previously enriched or co-cultured Lokiarchaeia, however, these Asgard archaea were found to possess large numbers of intracellular vesicles. While vesicles were only present in a small number of cells in late-stage cultures, they were common in early-stage cultures in cells that lacked protrusions and which were entering a period of rapid growth. Thus, *Ca.* Y. umbracryptum cells have a large surface to volume ratio at all stages of growth, and appear to be able to use vesicles and protrusions as interchangeable condition-specific membrane reservoirs. While an understanding of the triggers that cause cells taken from cultures at two different stages in their growth cycle to differ in their morphology and internal cell architecture will require a much more detailed future analysis, growth to stationary phase is likely accompanied by a host of changes in the chemical and physical extracellular environment, all of which may contribute to this effect – as has been shown to be the case for changes in the shape of other archaea^47,48.^

The majority of the vesicles present in *Ca.* Y. umbracryptum cells were lined on their luminal side by a proteinaceous inner coat that resembled the external surface of the plasma membrane (Fig. 4). Additionally, we observed a single instance of a pilus being present inside a non-spherical vesicle (Fig. 3d-f), implying that its contents were being trafficked either to or from the cell surface. Since vesicles appear to be formed and lost according to environmental conditions, we expect this traffic to flow in both directions. Note that vesicles in eukaryotic cells bud off from both the plasma membrane and the ER and Golgi membranes via similar but mechanistically distinct processes as they traffic cargo into or out of the cell, complicating direct comparisons with *Ca.* Y. umbracryptum cells that have a single continuous outer membrane.

We were unable catch intermediates in the process of vesicle fusion or fission. This would be expected if the process was fast. However, we cannot exclude the possibility that many of the vesicles were generated via a distinct archaeal-specific process. The ability of these cells to generate vesicles in an unusual way is also suggested by structures seen in these cells that are rarely seen in bacteria or eukaryotic cells, including vesicle membranes connected by three-way junctions (**Figs. 3d-e and Supplementary Fig. 10**), along with large numbers of vesicles within vesicles (**Figs. 3d-f and 4c-d**). At the same time, there were rare instances in which tomograms appeared to catch a new membrane compartment in the process of forming in a way that resembled the way multivesicular bodies are formed in eukaryotes^49^ - with cytoplasm and ribosomes becoming enclosed on the inside (**Figs. 3d-f**). In addition, we also observed a very few cases in which extracellular material was seen in a deep invagination in the external membrane of a cell in tomograms, and one case of a *Ca.* Y. umbracryptum cell ensheathing another cell in the mixed culture (**Supplementary Fig. 11a-c**). Moreover, while most vesicles were small, we observed rare instances in which cells had an internal compartment that contained dense material resembling abiotic material seen in the extracellular space (**Supplementary Fig. 11f**), which suggested the possibility that they may be able to undergo a process similar to phagocytosis. Thus, it seems possible that *Ca.* Y. umbracryptum can generate a range of internal vesicles of different types, perhaps via distinct processes.

The genome of *Ca.* Y. umbracryptum provides few clues as to nature of the molecular machinery underpinning vesicle formation. Despite the abundance of intracellular membrane-bound compartments observed in some cells, *Ca.* Y. umbracryptum has a small genome in which we were unable to identify homologues of Sec13/31, Clathrin, COP-II, all of which are known to function in membrane trafficking in eukaryotes^50,51^. Furthermore, *Ca.* Y. umbracryptum does not contain obvious homologues of dynamin, which drives membrane scission in eukaryotes, but which has been reported to be of bacterial origin^52^, nor did we observe any evidence of branched actin in the vicinity of vesicles or coat-like proteins in tomograms. In addition, *Ca.* Y. umbracryptum does not encode homologues of the proposed membrane trafficking associated complexes that are encoded in the genomes of other Asgard/Prometheoarchaeota^8,13^. Therefore, either the homologues of these proteins are far too divergent to be detected by standard bioinformatics or *Ca.* Y. umbracryptum cells use alternative machinery to generate internal vesicles. Nevertheless, the cells possess a large set of small GTPases^53^ and, despite the lack of classical endocytotic machinery in *Ca.* Y. umbracryptum (**Supplementary Fig. 3b-c**)^50^, we were able to identify multiple SPFH-family proteins in the genome that could plausibly form membrane-attached basket-like structures observed on the inner face of the plasma membrane (**Supplementary Fig. 7**). These proteins are ubiquitous across the tree of life^54,55^ and are implicated in a variety of functions, such as protease regulation, membrane signalling and, in case of eukaryotic flotillin, clathrin-independent endocytosis^43^. Whether or not they could contribute to membrane remodelling in *Ca.* Y. umbracryptum remains to be explored.

While an understanding of vesicle formation in this system will have to await improvements in our ability to perform live cell imaging of these small anaerobes at subcellular resolution, the discovery of *Ca.* Y. umbracryptum challenges established views of archaeal cell biology and the timeline of events leading to the origin of eukaryotes. Since abundant internal vesicles were not seen in recent cell biological studies of the Lokiarchaeia^7,29^, it is possible that intracellular membranes emerged in the Heimdallarchaeia which, under most phylogenetic analyses, are closer relatives of eukaryotes than Lokiarchaeia^8,13^,14 (**Fig. 4f**). Alternatively, it is possible that vesicles were missed in ultrastructural studies of Lokiarchaeal cells because they only form under specific conditions, for example, in cells that have yet to use up the membrane reservoir in the form of protrusions (depicted by a question mark in **Fig. 4f**).

Under either scenario, the capacity of *Ca.* Y. umbracryptum to generate and lose intracellular membrane-bound compartments in a way that depends on conditions supports a hypothesis in which the archaeal ancestor of eukaryotes had a dynamic endomembrane system prior to its association with the bacterial ancestor that gave rise to mitochondria. If the vesicles in *Ca.* Y. umbracryptum cells are used to exchange material with its environment to facilitate syntrophic partnerships, it is possible that an archaeal ancestor of eukaryotes engulfed the ancestor of mitochondria, perhaps aiding its ability to migrate into oxygen-rich environments^56^. In addition, if, as previously suggested^57^, the central cell body of cells that contains the genome can functionally separate transcription from sites of translation and protein activity at the cell periphery, these cells may also possess a proto-nuclear compartment^57^. In these and other ways, the cell biological analysis of different members of the Asgard archaea is closing the gulf that was long assumed to separate bacteria and archaea from eukaryotes.

### Etymology

*Candidatus* ‘Y. umbracryptum’. archaeum, from *archaea*, (Latin, Greek Origin); primitive. *Umbra* (Latin); shadow, *cryptum* (Latin, Greek Origin); vault. The name describes an archaeon containing hidden compartments.

### Sample site description

*Candidatus* ‘Y. umbracryptum’ was enriched from poorly lithified microbial mats collected from a tidal flat of a hypersaline pool in Shark Bay, Western Australia.

## Acknowledgements

B.B. and T.A.M.B. groups received support from the Medical Research Council, as part of United Kingdom Research and Innovation (also known as UK Research and Innovation) [Programme MC_UP_1201/31 to T.A.M.B] and [Programme MC_UP_1201/27 to B.B., which supported J.P. and I.R.]. B.B. also received support from the Wellcome Trust (222460/Z/21/Z), the Moore-Simons Project on the Origin of the Eukaryotic Cell, Simons Foundation 735929LPI and from the Gordon and Betty Moore Foundation’s Symbiosis in Aquatic Systems Initiative (9346). In addition, B.B. would like to thank the Allen Distinguished Investigator program, a Paul G. Allen Frontiers Group advised program of the Paul G. Allen Family Foundation for support for F.I.M. F.I.M. was initially supported by the Australian Government’s Research Training Program (RTP). T.A.M.B. would like to thank the Wellcome Trust (grant 225317/Z/22/Z) and the Lister Institute for Preventative Medicine for support. M.L. was supported by the Herchel Smith Fund PhD Scholarship. E.J.A-P. was supported by the Moore-Simons Project on the Origin of the Eukaryotic Cell, Simons Foundation grant 73592LPI to B.J.B., and a Simons Foundation investigator award LI-SIAME-00002001 to B.J.B. The authors also wish to thank Pieter Visscher, Tony Larkum, and Therese Morris for collecting the original microbial mat samples, as previously described^24,25.^ The authors acknowledge the support of Malgana rangers and coordinators in conducting the fieldwork. All fieldwork was conducted using valid permits issued by the Department of Biodiversity, Conservation and Attractions (DBCA), Western Australia. The original enrichment cultures were a kind gift from B.P.B to B.B. This study was also supported by the MRC Laboratory of Molecular Biology Electron Microscopy and Scientific Computing Facility. This research includes computations using the computational cluster Katana supported by Research Technology Services at UNSW Sydney. The authors thank Jan Löwe, Aurélien Roux, David Baum, Roni Levin-Konigsberg, Sean Munro and Gautam Dey for constructive suggestions on the manuscript. For the purpose of open access, the MRC Laboratory of Molecular Biology has applied a CC BY public copyright license to any Author Accepted Manuscript version arising. This preprint was typeset using a modified word template originally available at www.github.com/chrelli/bi-oRxiv-word-template.

## Author contributions

F.I.M. performed cell cultivation, genome assembly, live cell and HCR FISH imaging; A.v.K. carried out the cryo-ET experiments; J.P. and I.R. performed the ExM FISH experiments; F.I.M., A.v.K., T.A.M.B., B.B., M.L., J.P. and I.R. analysed experimental data; F.I.M., M.L., E.J.A-P. and B.J.B analysed the genomic data; F.I.M., A.v.K., T.A.M.B. and B.B. designed the research; B.P.B. oversaw fieldwork, initial cell cultivation and early project direction. F.I.M., A.v.K. T.A.M.B. and B.B. wrote the manuscript with the support of all authors. B.B., T.A.M.B., B.P.B. and B.J.B raised the funds to support the work.

## Data availability

Subtomogram averaging maps will be deposited in the Electron Microscopy Data Bank (EMDB). For more details see Supplementary Table 2.

## Correspondence and requests for materials

Correspondence and requests for materials should be addressed to the corresponding authors T.A.M.B. or B.B.

## Materials and Methods

### Growth of *Ca*. Y. umbracryptum

Poorly lithified smooth microbial mat samples collected in 2019 from the Nilemah tidal flats of Hamelin Pool in Shark Bay (26 27’336’’S, 114 05’762’’E) were used as the initial inoculum to establish enrichment cultures. Briefly, serum bottles containing 50 mL of salt solution reflective of Hamelin Pool seawater (50.7 g/L NaCl, 13.3 g/L MgSO_4_·7H_2_O, 7.23 g/L MgCl_2_·6H_2_O, 2.7 g/L CaCl_2_·2H_2_O, 1.4 g/L KCl)^58^ were supplemented with 0.05% (w/v) yeast extract, 0.05% (w/v) casamino acids and 100 µg/mL resazurin. Serum bottles were sparged with 80:20 N_2_:CO_2_ (v/v) gas and sealed with butyl rubber stoppers and crimp sealed. The solution was further reduced by the addition of 0.3 g/L L-cysteine HCl and 0.3 g/L Na_2_S·9H_2_O. Serum bottles were autoclaved prior to the addition of 1 mL/L SL10 trace element solution^59^, 3 mL/L Vitamin 10 Solution^60^ and the pH was adjusted to 7.4. To inoculate cultures, serum bottles were opened and continuously flushed with 80:20 N_2_:CO_2_ (v/v) gas and 2 g of homogenized microbial mat was added to each bottle. Each bottle was then resealed with butyl rubber stoppers and crimp sealed. The headspace was reestablished by flushing with 80:20 N_2_:CO_2_ (v/v) gas. Bottles were incubated at 37 °C for 90 days. For subsequent transfers over the next 270 days, 1 mL of enrichment culture was added to 49 mL of medium. The medium was unchanged bar the addition of 4 g/L microbial mat sample to the salt solution which was sterilized through repeated autoclaving and the final pH adjustment was performed by adjusting to pH 7.5 with a 10% (w/v) solution of Na_2_CO_3_. For subsequent transfers after 360 days post inoculation, media was adjusted to include 100 µg/mL (w/v) ampicillin, 50 µg/mL (w/v) kanamycin, and 100 µg/mL (w/v) streptomycin which was added after autoclaving and prior to pH adjustment. For three rounds of the enrichment, the media was additionally supplemented with 64 µg/mL of ciprofloxacin.

After a further four years of continuous cultivation under the previously described media, the media was adjusted to an alternative media herein referred to as Hamelin Pool Media (HPM, see Supplementary Table 3). HPM contained 50.7 g/L NaCl, 13.3 g/L MgSO_4_·7H_2_O, 7.23 g/L MgCl_2_·6H_2_O, 2.7 g/L CaCl_2_·2H_2_O, 1.4 g/L KCl and was supplemented with 0.05% (w/v) yeast extract and 0.05% (w/v) casein hydrolysate and 100 µg/mL resazurin in serum bottles. The bottles were again reduced by sparging with 80:20 N_2_:CO_2_ (v/v) gas and further reduced by the addition of 0.3 g/L L-cysteine HCl and 0.3 g/L Na_2_S·9H_2_O. The headspace was then established by flushing with 80:20 N_2_:CO_2_ (v/v) gas and setting final pressure to 0.5 bar. Bottles were then autoclaved. After autoclaving, 100 µg/mL (w/v) ampicillin, 50 µg/mL (w/v) kanamycin, and 100 µg/mL (w/v) streptomycin was added to each serum bottle along with 1 mL/L SL10 trace element solution, 3 mL/L V10 solution, 1 mL/L Se/W/Mo solution containing 0.021 g/L Na_2_SeO_3_, 0.037 g/L Na_2_WO_4_, 0.021g/L Na_2_MoO_4_ and 50 µL of 1 M KH_2_PO_4_ solution (pH 7.5). Finally, the pH was adjusted to pH 7.5 with a 10% (w/v) solution of Na_2_CO_3_. For subculturing in HPM, 5 mL of cell culture was added to 45 mL of HPM media and cultures were transferred every 60 days. The enrichment cultures were regularly checked for their abundance, both absolute and relative, through qPCR and amplicon sequencing, respectively. DNA was extracted from 1 mL of enrichment cultures using DNeasy PowerBiofilm Kits, according to manufacturer’s instructions. For qPCR, the copy number of 16S was determined by running extracted DNA on a ViiA 7 Real-Time PCR system using the PowerTrack SYBR Green Master Mix and the *Ca*. Y. umbracryptum 16S specific primers HEI245F (5’ - TCGGTAGGGGCTATGAGAGT-3’) and HEI416R (5’ - TTGCTGCCGGAG-TTTTACAC - 3’) against a standard curve of known template concentrations. For relative abundance determination, PCR was performed using the universal primer pair 926F (5’ - AAACTYAAAKGAATTGRCGG - 3’) and 1392wR (5’ - ACGGGCGGTGWGTRC - 3’). PCR products were cleaned using a Monarch PCR & DNA Cleanup Kit. A long-read sequencing library was prepared using a v14 library and sequenced by Oxford Nanopore Technologies. Quality checks were performed on fastq reads with FastQC and MultiQC^61,62.^ Adaptor sequences and chimeras were removed from fastq reads using Porechop^63^. Reads were filtered to meet quality standards using fastp^64,65.^ Read quality was then reevaluated using FastQC and MultiQC^61,62.^ Taxonomy was assigned to reads using Kraken2 using the SILVA SSU 138 database^66,67.^ Relative abundance of the Kraken 2 assigned reads was determined using Bracken^68^.

### Genome assembly, binning, and taxonomic placement

For characterization of the *Ca*. Y. umbracryptum genome, DNA was extracted from 5 mL of 90 day culture in triplicate using a potassium ethyl xanthogenate/sodium dodecyl sulfate (SDS) lysis buffer, as described previously^24^. DNA was purified from cell lysate using a phenol:chloroform:isoamyl alcohol extraction followed by an ethanol precipitation with the inclusion of glycogen. DNA quality and concentration was assessed using Xpose (Trinean, Belgium), NanoDrop (ThermoFisher Scientific, U.S.A) and Qubit (ThermoFisher Scientific, U.S.A) spectrophotometers. Short read library preparation was performed using a Nextera DNA Flex Library Prep Kit (Illumina, U.S.A). DNA was sequenced on a NovaSeq 6000 using a NovaSeq 6000 SP 2×150bp flowcell (Illumina, U.S.A). DNA sequencing was performed at the Ramaciotti Centre for Genomics (The University of New South Wales, Australia).

The quality of fastq reads was assessed with FastQC^61^ and low-quality bases were removed with Trimmomatic^69^. The resulting triplicate trimmed fastq reads were co-assembled into contigs using SPAdes v.3.15.5 with the - meta option^70,71.^ Assembled contigs were mapped against the unassembled DNA sequences using BWA-MEM^72^. Contigs shorter than 1000 bp were removed and resulting contigs were used to generate bins with Metabat v.2.12.1, CONCOCT v.1.0 and MaxBin2 v.2.2.3^73-75^. Bins generated were dereplicated and refined using Das Tool^76^. Genome quality checks for completeness and contamination were performed with CheckM2^77^. Barrnap v.0.9 was used to extract rRNAs^78^. tRNAs were extracted using tRNAscan-SE^79^. The nucleotide sequences of draft genomes were predicted with Prodigal v.2.6.3^80^.

Draft genomes were provided an initial taxonomic placement using GTDB-Tk against the GTDB database^26,81.^ A single Heimdallarchaeaceae MAG was detected as belonging to the Heimdallarchaeia order UBA460, family UBA460, genus novel, species novel (*syn*. order Heimdallarchaeales, family Heimdallarchaeaceae). Taxonomic placement was further validated by constructing a phylogenetic dataset of archaea for further phylogenetic analyses, including the target Heimdallarchaeaceae genome generated as part of this study. Briefly, a set of archaeal genomes was downloaded from GTDB on the 4th of August 2025 meeting the following criteria: CheckM2 completeness > 80%, CheckM2 contamination < 10%, and GTDB representative of species = true. The genome of highest completeness for each order was selected for DPANN archaea, Euryarchaeota and Thermoproteota, except in cases where orders contained well characterized representatives. The genome with the highest completeness for each Asgardarchaeota genus was included, and all Heimdallarchaeia above 90% completeness were included. Marker genes of the ar53 dataset were extracted with GTDB-tk identify^26^ and aligned with MAFFT v.7 using the L-INS-I iterative refinement method^82^. The resulting alignment was trimmed with BMGE with entropy estimation according to the BLOSUM30 matrix and a gas-rate cutoff of 50%^83^. BMGE trimmed alignments were further manually inspected and trimmed as required. Maximum-likelihood phylogenetic trees were constructed with IQ-TREE 2 with ultrafast bootstrap approximation (option - bb 10000) and model selection with ModelFinder Plus (selected as LG+F+R10)^84-86^. The resulting phylogenetic tree was visualized and annotated in iTOL^87^.

For the placement of the Heimdallarchaeaceae genome generated as part of this study in an archaeal tree including eukaryotes, the NM57 marker genes from a previously published taxonomic dataset were collected^8^. NM57 marker genes from the Heimdallarchaeaceae genome were annotated with eggNOG Mapper v.2.1.8 and aligned to reference markers with MAFFT v.7 using the L-INS-I iterative refinement method^82,88^. Trimming and phylogenetic tree calculation for the NM57 dataset was performed identically to the ar53 dataset (model LG+F+R10). The resulting phylogenetic tree was visualized and annotated with iTOL^87^.

### Genomic analysis

To functionally characterize the *Ca*. Y. umbracryptum genome, protein sequences were annotated against the CATH-Gene3D^89,90^, CDD^91^, HAMAP^92^, PANTHER^93^, Pfam^94^, PIRSF^95^, PRINTS^96^, PROSITE profiles^97^, PROSITE patterns^97^, SFLD^98^, SMART^99^, and SUPERFAMILY^100^ databases with InterProScan v.5.44-79.0^101^. The genome was additionally annotated against the eggNOG database with eggNOG Mapper v.2.1.8^88^ and KEGG database with Ghostkoala v.3.1^102^. The resulting InterProScan and Ghostkoala annotation were used to provide the annotation of ESPs and metabolic genes, respectively. Four SPFH-family protein candidates in the *Ca.* Y. umbracryptum genome were identified by Band-7/SPFH-domain annotation (PANTHER: IPR043202; SMART and Pfam: IPR001107; SUPERFAMILY: IPR036013; Pfam and CDD: IPR033880). AlphaFold3^103^ modelling of the candidates revealed two putative proteins with an elongated alpha-helix following the domains SPFH1 and 2, typical for basket-forming SPFH family members (NODE_169_length_248565_cov_475.496599_54 and NODE_955_length_102699_cov_479.414219_56, named CC1-SPFH A and CC1-SPFH B respectively). Representatives of basket-forming proteins from other organisms were chosen from the literature and by taxonomic search of the Band-7 domain containing proteins annotated in InterPro (IPR001107)^104^. Detailed domain annotation was performed based on AlphaFold3^103^ structural prediction of monomeric proteins. All structures shown were rendered using ChimeraX^105^.

### Light microscopy

Cells from late-stage (stationary phase) cultures in duplicates were allowed to settle onto polyethyleneimine-coated glass imaging chambers in an anaerobic chamber before a brief centrifugation at 1000 rcf (relative centrifugal force). Cells were then stained with Mitotracker Red CMXRos (200 nM, Invitrogen) and Hoechst 33342 (40 µM, Invitrogen) to label the membranes and DNA, respectively. Wells were overlaid with mineral oil to minimise oxygen exposure, and cells were immediately imaged on a SoRa spinning disk confocal microscope to avoid oxygen-induced toxicity over time. Images were collected with a pixel size of 27.5 nm^2^. An identical methodology was used to image cells in early-stage cultures, which had yet to reach stationary phase (see Fig. 1c).

### HCR FISH microscopy

For the identification of *Ca*. Y. umbracryptum in enrichment cultures by light microscopy, cells of early-stage and late-stage cultures (at times indicated in Fig. 1c) in duplicate were allowed to settle onto 1% polyethyleneimine-coated glass imaging chambers in an anaerobic chamber prior to centrifugation at 1000 rcf. Cells were then fixed with 1.6 % formaldehyde at 4 °C for 15 minutes. After formaldehyde fixation, cells were washed three times with the anaerobic Hamelin Pool Salt (HPS) buffer. HPS buffer contained 50.7 g/L NaCl, 13.3 g/L MgSO_4_·7H_2_O, 7.23 g/L MgCl_2_·6H_2_O, 2.7 g/L CaCl_2_·2H_2_O, 1.4 g/L KCl, which was sparged with 80:20 N_2_:CO_2_ (v/v) gas, and adjusted to pH 7.5 with 10 % (w/v) Na_2_CO_3_. Cells were then permeabilised with 0.01 % (w/v) SDS in HPS for 15 minutes at room temperature. To specifically tag *Ca*. Y. umbracryptum 16S rRNA fluorescently, Hybridised Chain Reaction Fluorescent in situ Hybridisation (HCR-FISH) probes were designed for the target organism by Molecular Instruments (U.S.A). Cells were pre-hybridised using HCR HiFi Probe Hybridization Buffer for 10 minutes at 37 °C. Cells were hybridized with 2 µL HCR HiFi Probe in 100 µL hybridisation buffer at 37 °C overnight. Cells were then washed four times with HCR HiFi Probe wash buffer at 37 °C for 15 minutes, and incubated with 200 µL of HCR Gold Amplification buffer at room temperature for 30 minutes. Amplified hairpins were heated to 95 °C for 90 seconds before cooling to room temperature in the dark for 30 minutes. Hairpins were then added to HCR Gold Amplifier Buffer (2 µL hairpin to 100 µL amplifier buffer), and cells incubated in the dark with the hairpin HCR Gold Amplifier Buffer solution at room temperature overnight. Cells were then washed with HCR Gold Amplifier Wash Buffer four times for 15 minutes at room temperature. Finally, cells were stained with NHS ester 488 in phosphate buffered saline (PBS) for at least one hour at room temperature prior to three washes with PBS, overlaid with 200 µL of PBS, and imaged on a SoRa spinning disk confocal microscope (pixel size 27.5 nm^2^).

### HCR FISH Expansion microscopy

We adapted a previously described protocol^31^ for our experiments. Briefly, cells were washed three times for 15 minutes each with sodium bicarbonate (100 mM) before treatment with Glycidyl methacrylate (GMA, 0.08 %) dissolved in sodium bicarbonate (100 mM) for 3 hours at room temperature. Cells were then cross-linked overnight in freshly prepared 1 % acrylamide/0.7 % formaldehyde solution in PBS. The following day cells were washed once with PBS before the addition of gelation solution (19 % (w/w) sodium acrylate, 10% (w/w) acrylamide, 0.1 % (w/w) bis-acrylamide in 10x PBS), each 18 µL aliquot was activated with 1 µL 10 % (w/v) ammonium persulfate (APS) and 1 µL tetramethylethylenediamine (TEMED) and mixed by vortexing to make two gels of 9 µL solution each. The gels were left on ice for 5 minutes before transfer to a humidified chamber which was placed at 37 °C for at least one hour. Following gelation, the cells were denatured at 95 °C with denaturation buffer (50 mM TRIS pH 8, 200 mM NaCl and 200 mM SDS in Millipore water). Following denaturation, gels were left to cool to room temperature before washing twice with PBS and then four successive 30-minute expansion steps with Millipore water at room temperature on a shaker. The water was exchanged once more and left at 4 °C overnight to fully expand. After overnight expansion, the full-size gels were cut into four quarters and shrunk by two 15-minute PBS washes on a shaker before storage in PBS with 0.02 % (w/v) sodium azide at 4 °C.

HCR-FISH was used to label rRNA for light microscopy. First, the HCR-HiFi Probe Hybridization Buffer was pre-heated to 37 °C. The expansion gels were then pre-hybridized in 200 µL of HiFi Probe Hybridization Buffer at 37 °C for 30 minutes. The pre-hybridisation solution was then removed, and 100 µL of probe hybridisation solution containing the HCR probes (2 pM) was added. The samples were then left to hybridise overnight at 37 °C without agitation. The samples were then washed in 500 µL of HCR HiFi Probe Wash Buffer (4 x 15 minute) at 37 °C. Following this, the samples were pre-amplified for 30 minutes at room temperature in 200 µL of HCR Gold Amplifier Buffer. 3 µL of hairpin-1 and 3 µL of hairpin-2 per sample were heated for 90 seconds at 95 °C and then cooled at room temperature in the dark. The amplifier solution was then made by combining the hairpin-1 and hairpin-2 into 100 µL (per sample) of HCR Gold Amplification Buffer at room temperature. The pre-amplification solution was then removed and replaced with the amplifier solution containing hairpins. The samples were again left to incubate overnight at room temperature in the dark for amplification. The amplifier solution was then removed, and excess hairpins were removed from the sample by washing 4 times for 15 min with 500 µL of HCR Gold Amplifier Wash Buffer at room temperature. Samples were then washed with PBS. Cells embedded in the gels were stained with NHS ester 647 or Bodipy in PBS (1:750) at room temperature for at least an hour before two washes with PBS followed by 3 × 20 min expansion steps with Millipore water. Gels were then imaged on the same day using the microscopy setup described above.

### Cryo-ET sample preparation

Cryo-ET grid preparation was performed either completely anaerobically within an anaerobic glove box using a manual plunger or with a Leica EM GP2 under ambient conditions. Within the anaerobic chamber, rubber caps of serum bottles were sterilised with 70 % (v/v) ethanol and ∼100 µL of sample were aseptically removed using a metal syringe. Prior to accessing cells, syringes were coated with 2 mg/mL Bovine Serum Albumin (BSA) (to minimise sample loss due sticking to the needle), formulated in 0.9% saline solution and then washed with 70 % (v/v) ethanol before being allowed to dry in an anaerobic chamber. Next, 3.5 µL of the specimen were directly applied to a freshly glow discharged Quantifoil R2/2 Au 200 mesh grid, blotted for 4-5 s and plunge-frozen into liquid ethane with a manual plunger. Vitrified grids were then transferred under liquid nitrogen, clipped and stored until cryo-ET data collection was performed. Similarly for ambient condition grid preparation, samples were aseptically removed from serum bottles and then 3.5 µL of the specimen was directly applied to a freshly glow discharged Quantifoil R2/2 Au 200 mesh grid, blotted for 4-5 s and plunge-frozen into liquid ethane with a Leica EM GP2, while the blotting chamber was maintained at 100% humidity at 18 ºC. Grids were clipped and stored under liquid nitrogen until cryo-ET data collection was performed.

### Cryo-ET data collection

Cryo-ET data was collected on a either Titan Krios G3 equipped with a K3 Gatan detector behind a Gatan Imaging Filter (20 eV slit width) or a Titan Krios G4 equipped with a Falcon 4i camera behind a Selectris-X Energy filter (10 eV slit width) using the SerialEM software^106^ and a dose symmetric tilting scheme^107^ at 300 kV. Three tilt series datasets for late-log phase *Ca.* Y. umbracryptum cells were collected at a pixel size 3.01 Å with a defocus range of-4 to-8 µm, ±60° oscillation, 3º increments with a total dose of ∼140 e^−^/Å^2^ using the Titan Krios G4 (see Supplementary Table 2). Another biological replicate for late-log phase *Ca.* Y. umbracryptum cells were collected using the Titan Krios G3 at a pixel size 2.13 Å with a defocus range of-5 to-8 µm, ±60° oscillation, 2º increments with a total dose of ∼120 e^−^/Å^2^. All early-log phase *Ca.* Y. umbracryptum cells and replicates were collected using the Titan Krios G3 at a pixel size 2.13 Å with a defocus range of-5 to-8 µm, ±60° oscillation, 2º increments with a total dose of ∼120-122 e^−^/Å^2^.

### Cryo-ET data analysis and subtomogram averaging

Cryo-ET data processing was performed using RELION5.0^32^. After tilt series import, tilt-series movies were motion-corrected and split into odd and even images with MotionCor2^108^ implemented in RELION-5.0. Tilt-series alignment and initial contrast transfer functions (CTFs) estimation were performed using ARETOMO2^109^. Tomograms were generated using RELION5.0 with options (--Fc --ctf –pre_weight --SNR 100) and denoised with Cryo-CARE^110,111^. Ribosomes were initially picked with pytom-match-pick^112^ using EMDB-10223^34^ as a reference and manually inspected using the ArtiaX plugin^113^ implemented in UCSF ChimeraX^105^. Subtomogram averaging was performed using RELION5.0 for multiple cycles of 2D-tilt-image extraction, realignment and CTF-optimisation as described previously^33^ (see Supplementary Table 2 and Supplementary Fig. 6 and 9). Figure panels containing cryo-ET images were prepared using IMOD^114^, Fiji^115^ and UCSF ChimeraX^105^ using the ArtiaX plugin^113^.

### Semantic and membrane segmentation

Semantic segmentation was performed using Dragonfly (2024.1.0.1627)^116^. Briefly, Cryo-CARE denoised tomograms were converted into float32 TIFF files using the newstack function of the IMOD^114^ package and then imported into Dragonfly with flip around the Y-axis option. Features (membrane, cell surface, filaments, pili, vesicles and inorganic granules) were manually annotated using a Wacom Cintiq Pro 16 pen display on three to five slices per tomogram as recently described^117^. Remaining un-labelled voxels were assigned to a background class and then used to train a semantic segmentation model with a 2.5D U-Net^118^ architecture containing 5-slices and a depth level of 5. Training was performed with a patch size of 128 over 100 epochs using Categorical Cross-Entropy loss function with the Adadelta optimisation algorithm^119^. All other training parameters were kept as default within Dragonfly. After training, segmentation was performed within Dragonfly, wrongly assigned voxels were manually cleaned using the process island function and then rendered for visualisation using UCSF ChimeraX^105^.

Membrane segmentation was performed using Membrain v2^120^ with default parameters. For vesicle quantification, the Membrain segmentation was imported into Dragonfly, and each vesicle was manually segmented as a separate feature. Highly elliptical vesicles were excluded from further analysis and elongated Membrain segmentation in the Z-direction were not annotated due to the missing wedge. All features/vesicles were exported as multi-ROI grey-value TIFF images with each integer value corresponding to one features/vesicles. Vesicle diameter and category (single, outer-filled or inner) were quantified using a custom python script. Briefly, 1) for each vesicle a sphere was fitted to binarized density of the vesicle, 2) the radius and centroid of the fitted sphere was saved, 3) and the category was computed by comparing the centroid and radius of each vesicle to all other vesicles of the same tomogram. If the centroid of a smaller radius vesicle fell within the centroid and radius of another vesicles it was assigned as an inner vesicle, or vice-versa as an outer vesicle. If neither condition is met it was assigned as single vesicle. All fits and assignment were manually inspected using a ChimeraX marker (.cmm) file containing the centroid and radius for each vesicle. Data was plotted using seaborn^121^.

### Two-dimensional class averages

Two-dimensional class averages of filaments and membranes were generated by converting binarized filament, membrane or vesicles segmentations generated above, into coordinates with a 10-voxels radial cutoff. Coordinates were then imported into RELION5.0 and classical 3D-subtomograms were extracted from CTF-corrected 4x down-sampled tomograms as implemented in RELION5.0. The central three slices were then Z-projected using a custom python script and used for 2D classification implemented in RELION5.0.

