## Supplementary figures and tables for "An Asgard archaeon with internal membrane compartments"

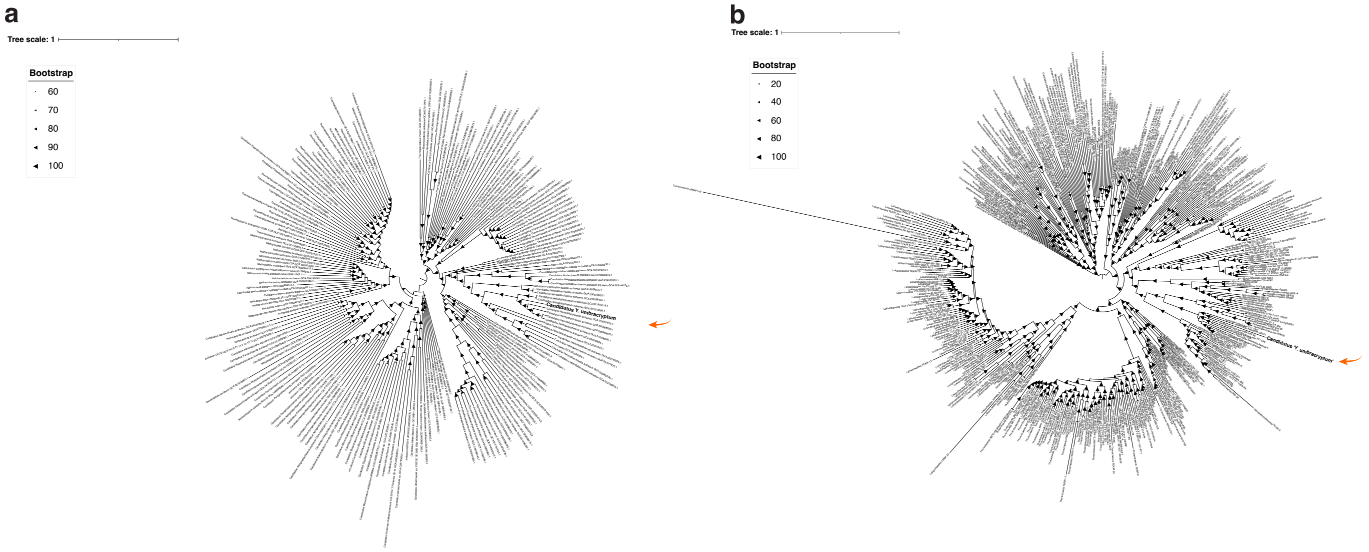


**Supplementary Figure 1: Phylogenetic placement of *Ca.* Y. umbracryptum.**
**(a)** Uncollapsed maximum-likelihood phylogenetic tree of *Ca.* Y. umbracryptum within the archaea. The phylogenetic tree was built using ar53 marker genes, as depicted in Fig. 1d. The tree was calculated under LG+F+R10 with ultrafast bootstrap approximation. **(b)** Uncollapsed maximum-likelihood phylogenetic placement of *Ca.* Y. umbracryptum within the archaea including the placement of Eukaryotes. The phylogenetic tree was built using the NM57 marker genes, as depicted in Fig. 1e. The tree was calculated under LG+F+R10 using ultrafast bootstrap approximation.

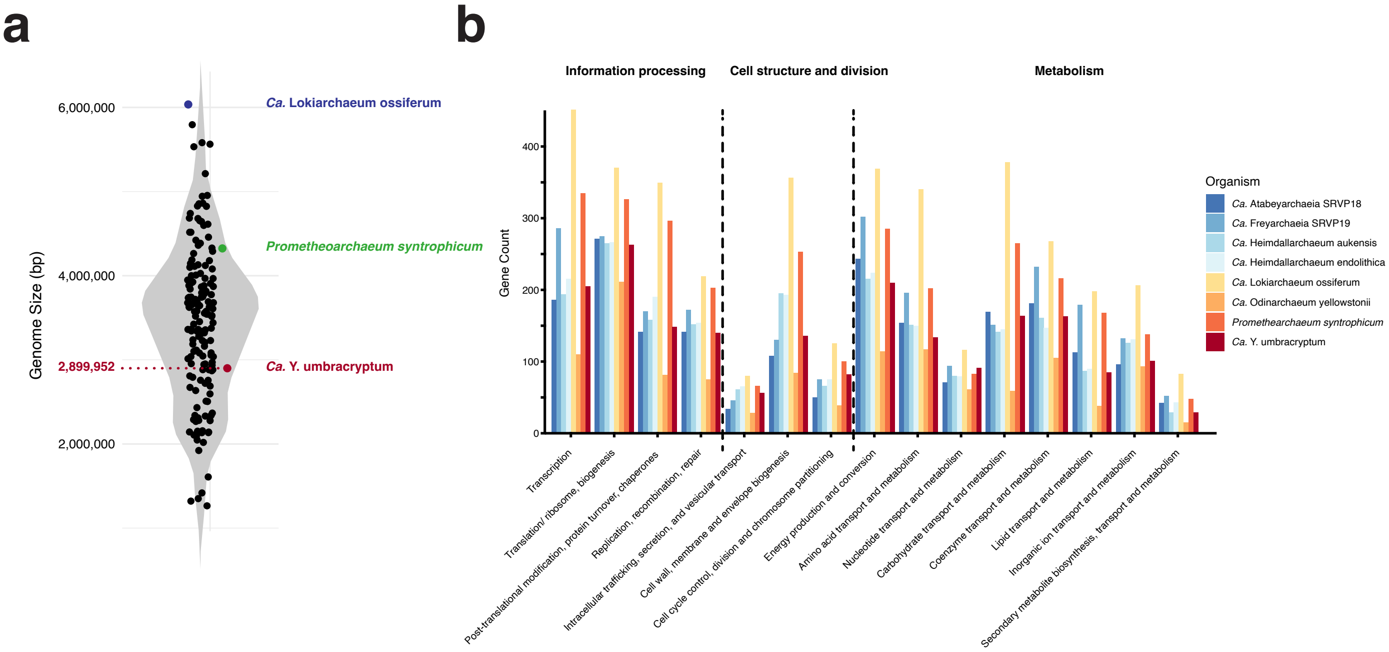


**Supplementary Figure 2: Genome size and distribution.**
**(a)** Genome size of *Ca.* Y. umbracryptum relative to other Asgard archaea. The genome was estimated to be >90% complete (see Methods) and genome size is illustrated relative to complete Asgard archaeal genomes downloaded from GTDB[81]. **(b)** Genome analysis showing the proportion of different gene families. *Ca.* Y. umbracryptum encodes a reduced number of genes related to various processes, including information processing, cell structure and division, and metabolism relative to *P. syntrophicum*, *Ca.* L. ossiferum, but was comparable to other Heimdallarchaeaceae.

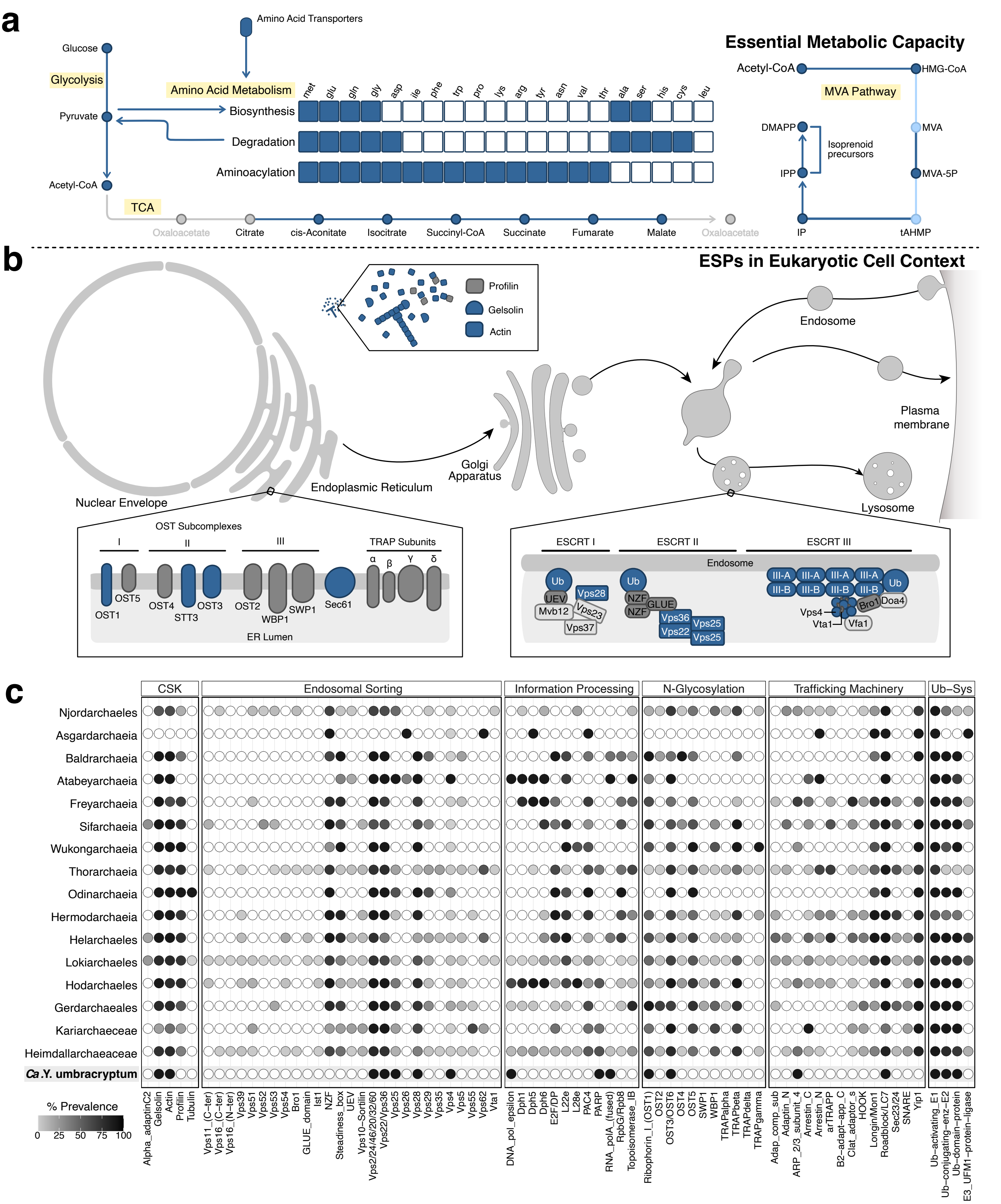


**Supplementary Figure 3: Reduced metabolism and ESPs in Eukaryotic context of *Ca.* Y. umbracryptum.**
**(a)** Diagram shows the essential metabolic capacity of *Ca.* Y. umbracryptum. Genes encoding metabolic enzymes were annotated with Interproscan v.5. and GhostKoala v.3.1, and results were verified manually. Blue denotes the presence of a gene, the ability to perform a given reaction, or produce a specific metabolic product. Specific metabolic pathways are highlighted in light yellow for reference. Abbreviations are MVA: mevalonate; TCA: tricarboxylic acid cycle. **(b)** Cartoon shows proteins identified in the *Ca.* Y. umbracryptum genome with clear eukaryotic counterparts within the context of a eukaryotic cell context. Blue denotes the presence of a gene; grey denotes genes that were previously identified in another Asgard genome. **(c)** Table shows the presence or absence of genes previously identified with close eukaryotic homologues across the Asgard archaea, with *Ca.* Y. umbracryptum highlighted in the bottom row. Prevalence of a gene within a given taxonomic group is defined by greyscale (bottom left). Values are based on recent publications' searches of Asgard genomes[8,122]. The analysis of ESPs in Asgardarchaeota clades, other than Atabeyarchaeia, has

been described previously[8]. ESPs within Atabeyarchaeia were identified using Pfam (v.34) and Interproscan (v5.61-93.0-64). These methods were validated with analysis of the arCOGs and AsCOGS in complete genomes of Atabeyarchaeia[122]. ESPs within *Ca.* Y. umbracryptum were identified using Interproscan v.5. Abbreviations: Ub-Sys: ubiquitin system; CSK: cytoskeleton.[8,122] ESCRT-III proteins fall into two classes, III-A and III-B[4,18,19].

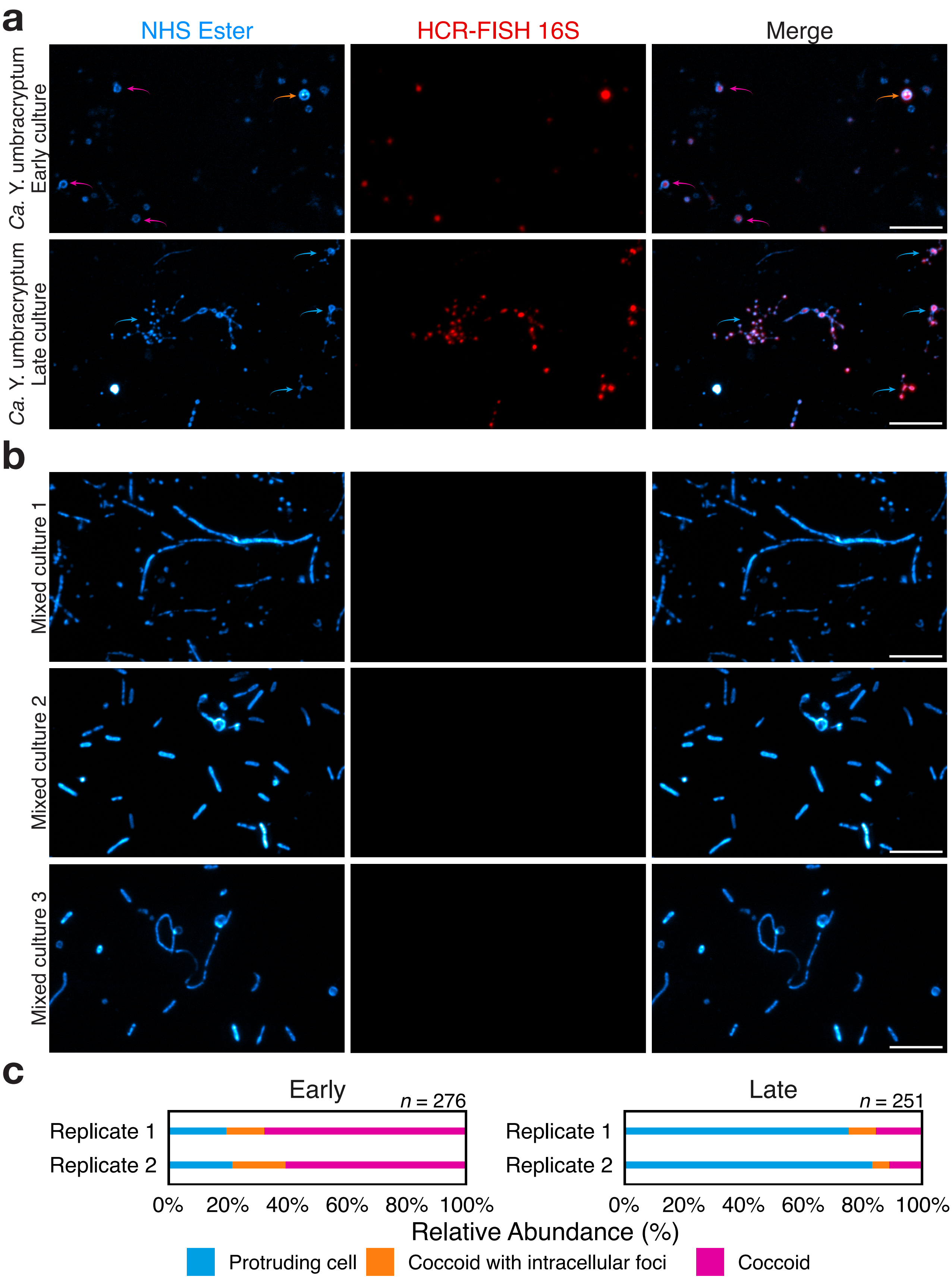


**Supplementary Figure 4: Light microscopy HCR.**
**(a)** Images show overviews of early and late-stage cultures of *Ca.* Y. umbracryptum labelled with specific HCR-FISH probes. Coccoid cells are indicated with pink arrows; coccoid cells with intracellular foci are indicated with orange arrows; cells with protrusions are indicated with blue arrows. Scale bars: 5 $\mu$m. **(b)** Images show controls for specificity of the *Ca.* Y. umbracryptum HCR-FISH probes in three biological replicates in cultures that have lost their population of *Ca.* Y. umbracryptum cells under identical imaging conditions. Cells lack an HCR-FISH positive signal. Scale bars: 5 $\mu$m. **(c)** Quantification of the morphology of HCR-FISH positive cells in early and late-stage cultures. Cells in the different morphological classes were counted across two biological replicates for both early and late-stage cultures.

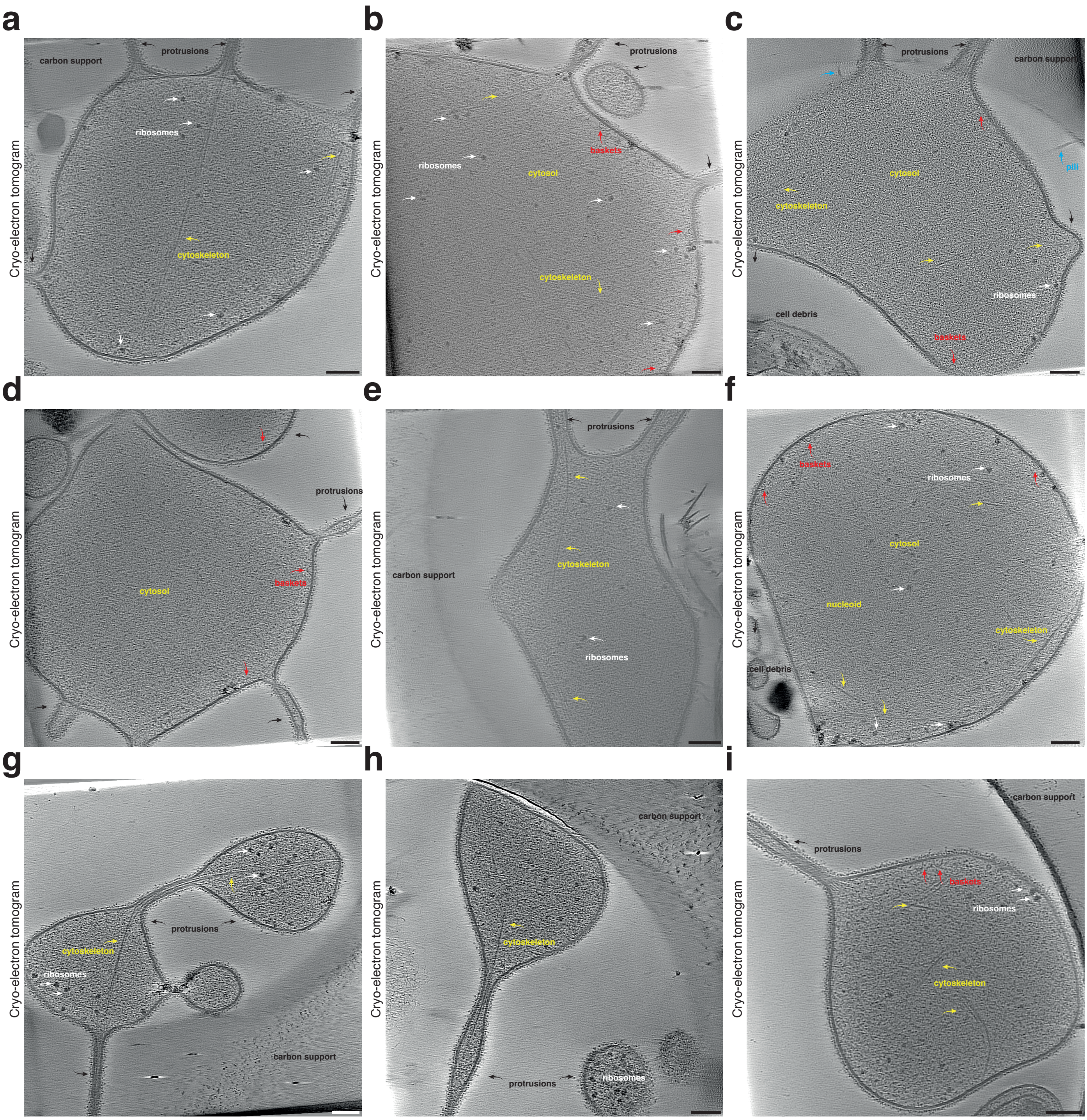


**Supplementary Figure 5: Cryo-ET of stationary phase *Ca.* Y. umbracryptum cells.**
**(a-i)** Cryo-ET slices through cells and protrusions of *Ca.* Y. umbracryptum with ribosomes (white arrows) and cytoskeletal filaments (yellow arrows) marked. Scale bars: 100 nm. Tomograms: $n$=135, from two biological replicates.

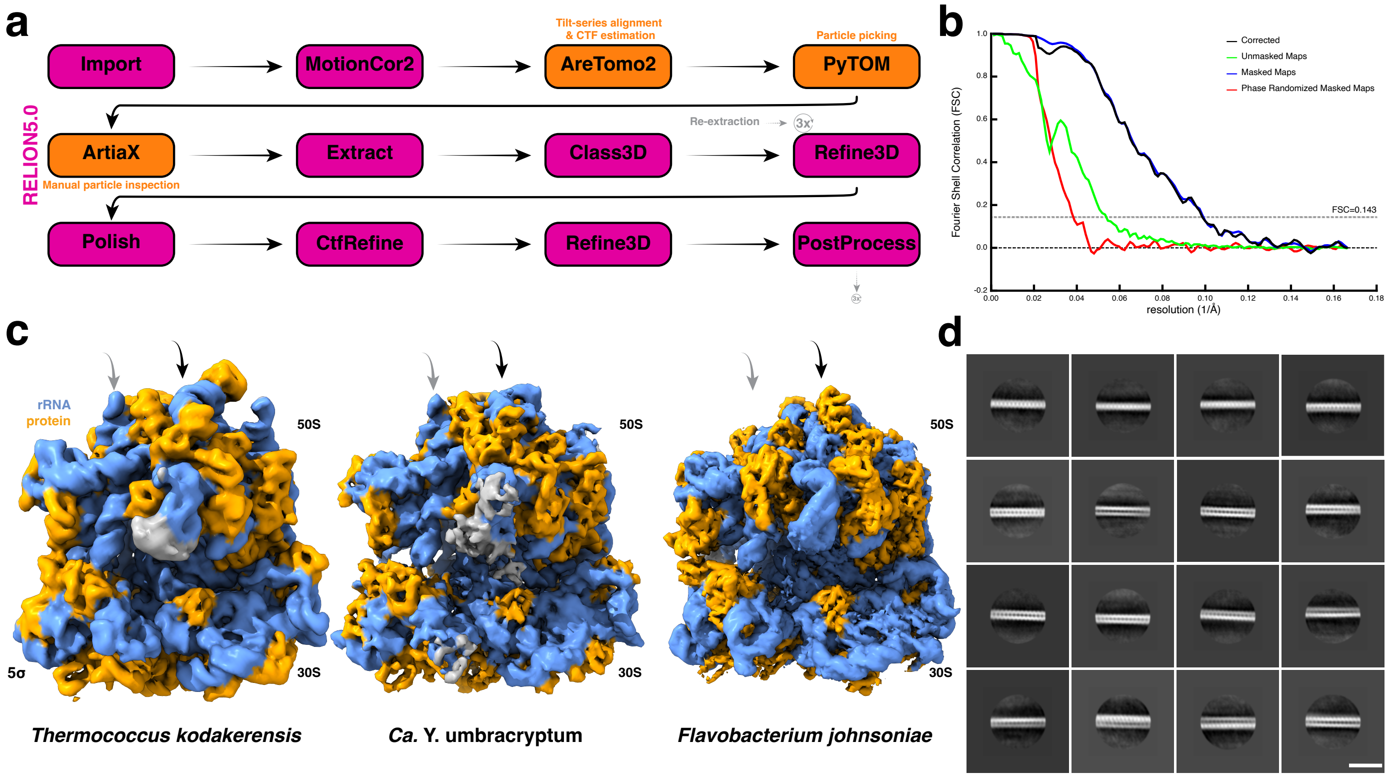


**Supplementary Figure 6: Subtomogram averaging of late stage ribosome and actin.**
**(a)** Subtomogram averaging workflow using RELION5.0[32] (magenta). After initial particle extraction, classification and refinement, three rounds of Bayesian polishing, CtfRefine and Refine3D were performed to obtain a final reconstruction with an estimated resolution of ~10 Å (also see Supplementary Table 2). **(b)** Fourier Shell Correlation (FSC) curves of the subtomogram averaging reconstruction between two independent half maps. **(c)** Comparison of the 70S ribosome structure of an archaeon (*T. kodakerensis*[34] -left; EMD-10223), *Ca.* Y. umbracryptum (center), and a bacterium (*Flavobacterium johnsoniae*[35] – right; EMD-22345). The cross-correlation values of the archaeal and bacterial ribosome maps are 0.93 and 0.90 when compared to the *Ca.* Y. umbracryptum ribosome, suggesting that the putative *Ca.* Y. umbracryptum ribosome is structurally closer to an archaeal, rather than a bacterial ribosome. The *F. johnsoniae* ribosome was chosen as representative of the class Bacteroidia (following GTDB taxonomy[81]) forming the highest abundant bacterial member in the culture (see Fig. 1b, left). All cryo-EM maps are filtered to 10 Å and displayed at a contour level of 5 sigma above the mean. **(d)** Two-dimensional class average of cytoskeletal filaments with a repeat of ~5.9 nm, seen inside *Ca.* Y. umbracryptum cells, the only species with a close actin homologue encoded on their genome. Scale bar: 50 nm.

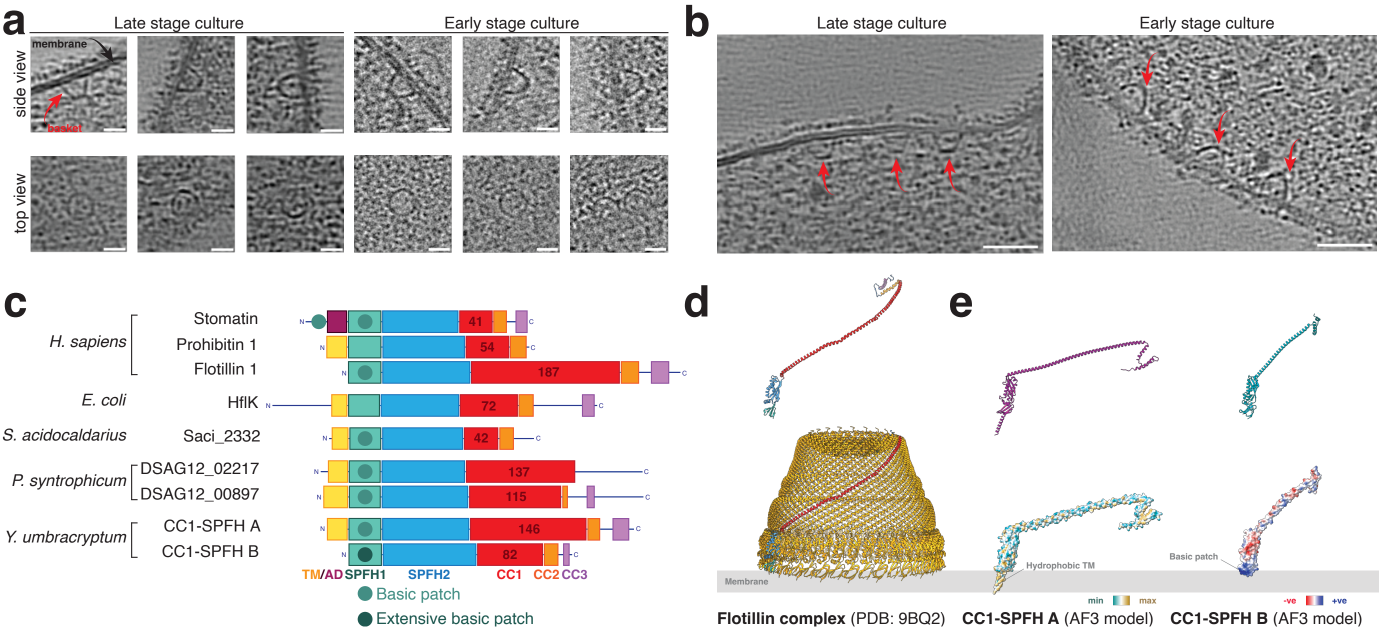


**Supplementary Figure 7: Cryo-ET of basket-like structures in *Ca.* Y. umbracryptum cells.**
**(a)** Cryo-ET slices of basket-like structures associated with the membranes in stationary/late stage and early-log phase cultures. Scale bars: 20 nm. **(b)** Examples of clusters of basket-like structures on the membrane in late-stage and early-stage cultures. Scale bars: 50 nm. **(c)** Cartoon representation of putatively basket-forming SPFH family proteins across the tree of life. Domain annotation: TM - transmembrane helix, AD - amphipathic domain, CC1-3 alpha-helical domains. **(d)** Human Flot-1 monomer (top) colored by domain composition and 44-subunit Flot1/2 complex (bottom). PDB: 9BQ2[42]. **(e)** AlphaFold3[103] models of CC1-SPFH A and CC1-SPFH B proteins from *Ca.* Y. umbracryptum genome (top). Bottom-left: surface lipophilicity of CC1-SPFH A suggests presence of transmembrane helix. Bottom-right: surface charge of CC1-SPFH B suggests the presence of a basic patch which could interact with a membrane composed of lipids with negatively charged headgroups.

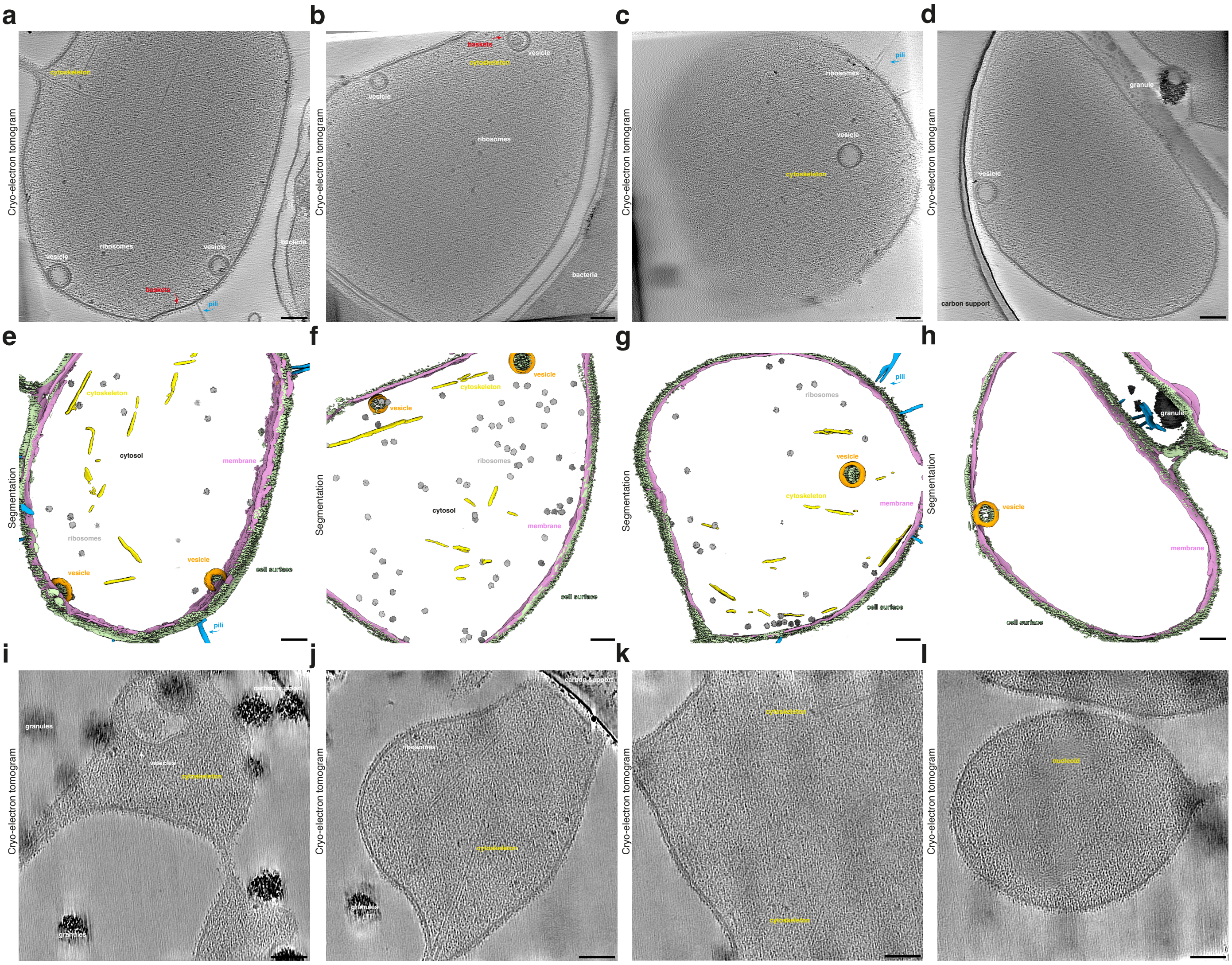


**Supplementary Figure 8: Cryo-ET of vesicle containing stationary phase *Ca.* Y. umbracryptum cells.**
**(a-d)** Cryo-ET slices through stationary phase *Ca.* Y. umbracryptum cells with intracellular vesicles. Scale bars: 100 nm. **e-h**, Semantic segmentations of tomograms shown in panels a-d) depicting the cell membrane (pink), cell-surface proteins (pale green), pili (blue), ribosomes (grey), cytoskeletal filaments (yellow) and intracellular vesicles (orange). One of the vesicles depicted in panel b) is coated by a basket-like structure (red – see Supplementary Fig. 7). Scale bars: 100 nm. Tomograms: *n*=21, from two biological replicates. **(i-l)** Cryo-ET slices through stationary phase *Ca.* Y. umbracryptum cells vitrified under strict anaerobic conditions within an anaerobic chamber. The cells display similar cellular features including intracellular membranes i), cytoskeletal filaments j-k), dense nucleoid l). Scale bars: 100 nm. Tomograms: *n*=25, from one biological replicate..

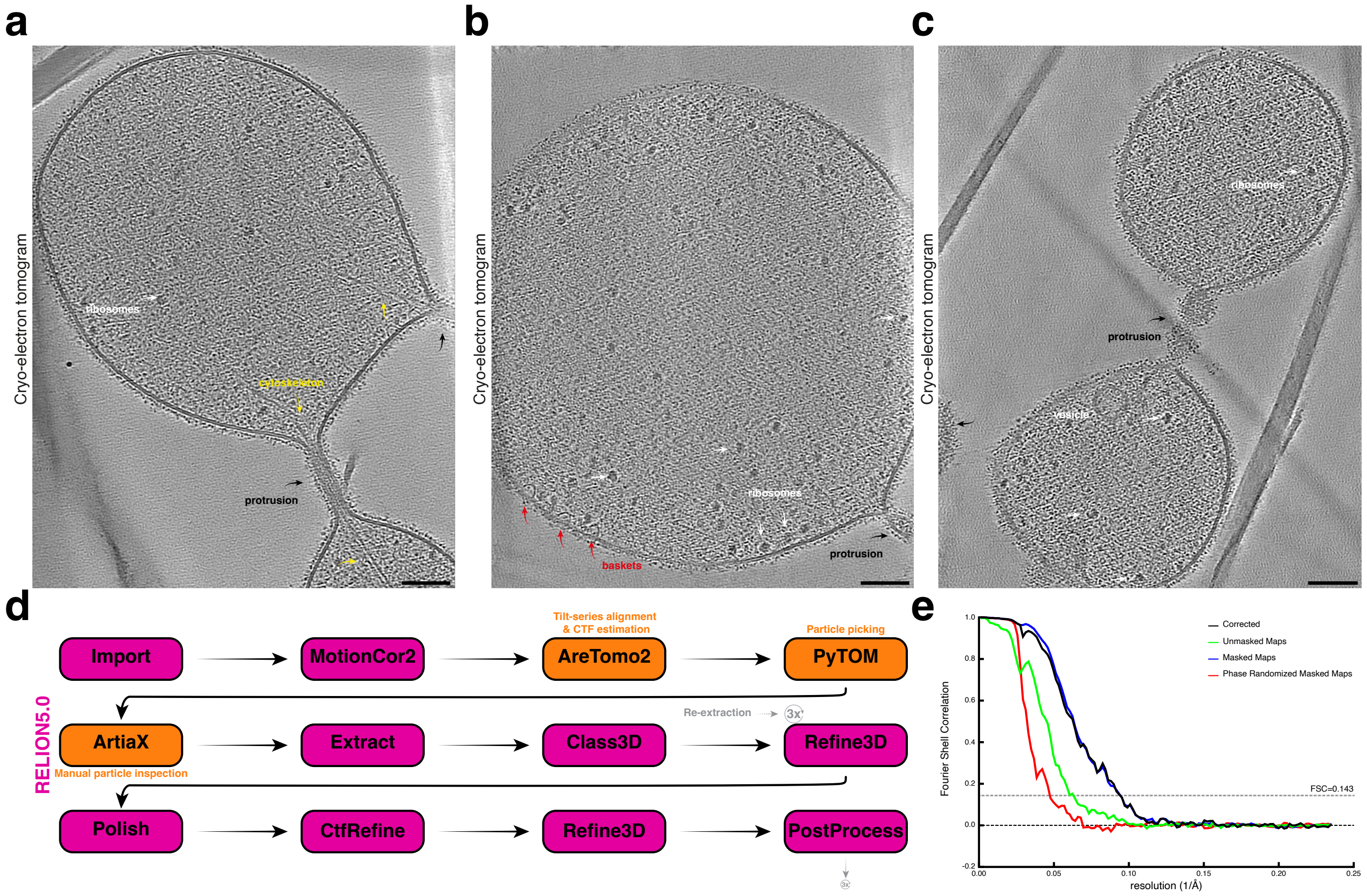


**Supplementary Figure 9: Cryo-ET of early-log *Ca.* Y. umbracryptum and ribosome STA.**
**(a-c)** Cryo-ET slices through early-log phase *Ca.* Y. umbracryptum cells containing protrusions with ribosomes (white arrows) and cytoskeletal filaments (yellow arrows) marked. Scale bars: 100 nm. (also see Supplementary Fig. 7b - right). **(d)** Subtomogram averaging workflow using RELION5.0[32] (magenta). After initial particle extraction, classification and refinement, three rounds of Bayesian polishing, CtfRefine and Refine3D were performed to obtain a final reconstruction with an estimated resolution of ~10.7 Å (also see Supplementary Table 2). **(e)** Fourier Shell Correlation (FSC) curves of the subtomogram averaging reconstruction between two independent half maps.

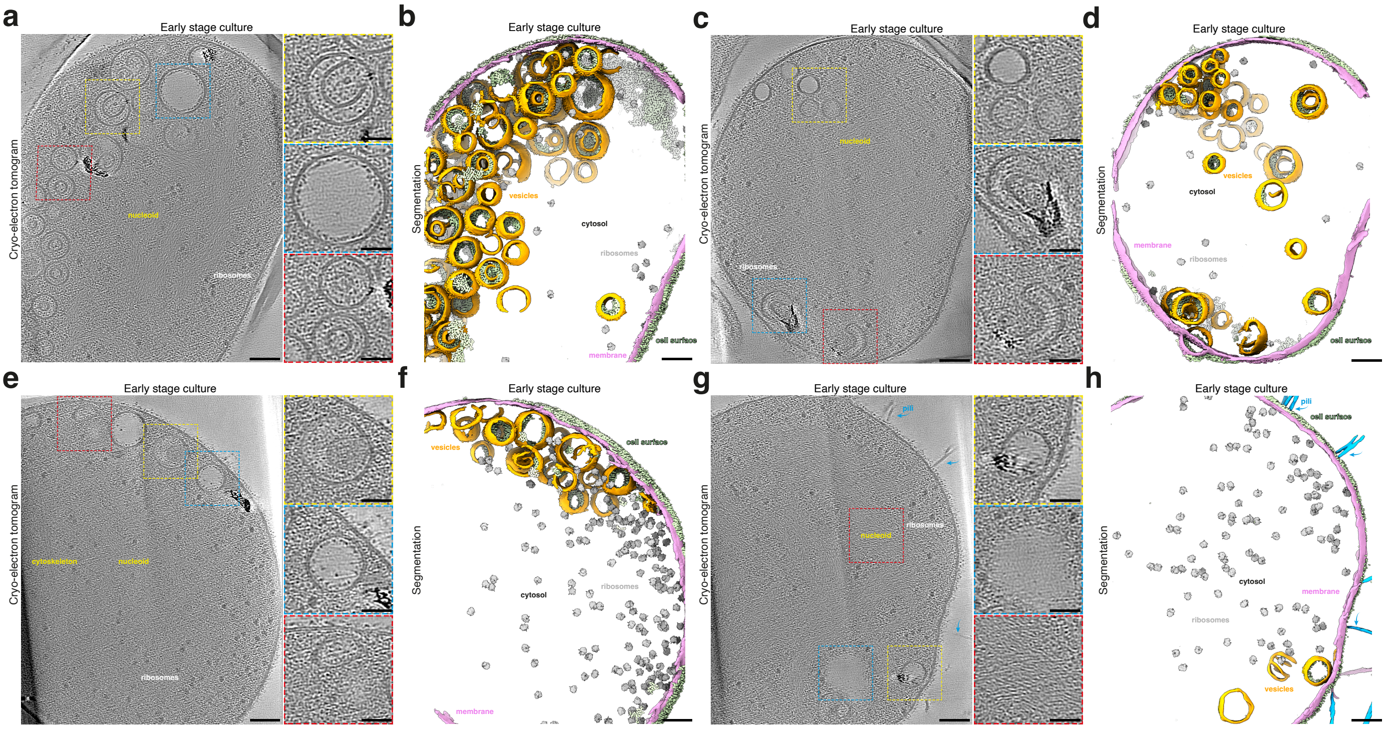


**Supplementary Figure 10: Cryo-ET of vesicle containing early-log phase *Ca.* Y. umbracryptum cells.**
**(a,c,e,g)** Cryo-ET slices through early-log phase *Ca.* Y. umbracryptum cells with intracellular vesicles. Scale bars: 100 nm. **(b,d,f,h)** Semantic segmentations of tomograms shown in panels a-d) depicting the cell membrane (pink), cell-surface proteins (pale green), pili (blue), ribosomes (grey), cytoskeletal filaments (yellow) and intracellular vesicles (orange). Scale bars: 100 nm. Close-up scale bars: 50 nm. Tomograms: *n*=35, from two biological replicates.

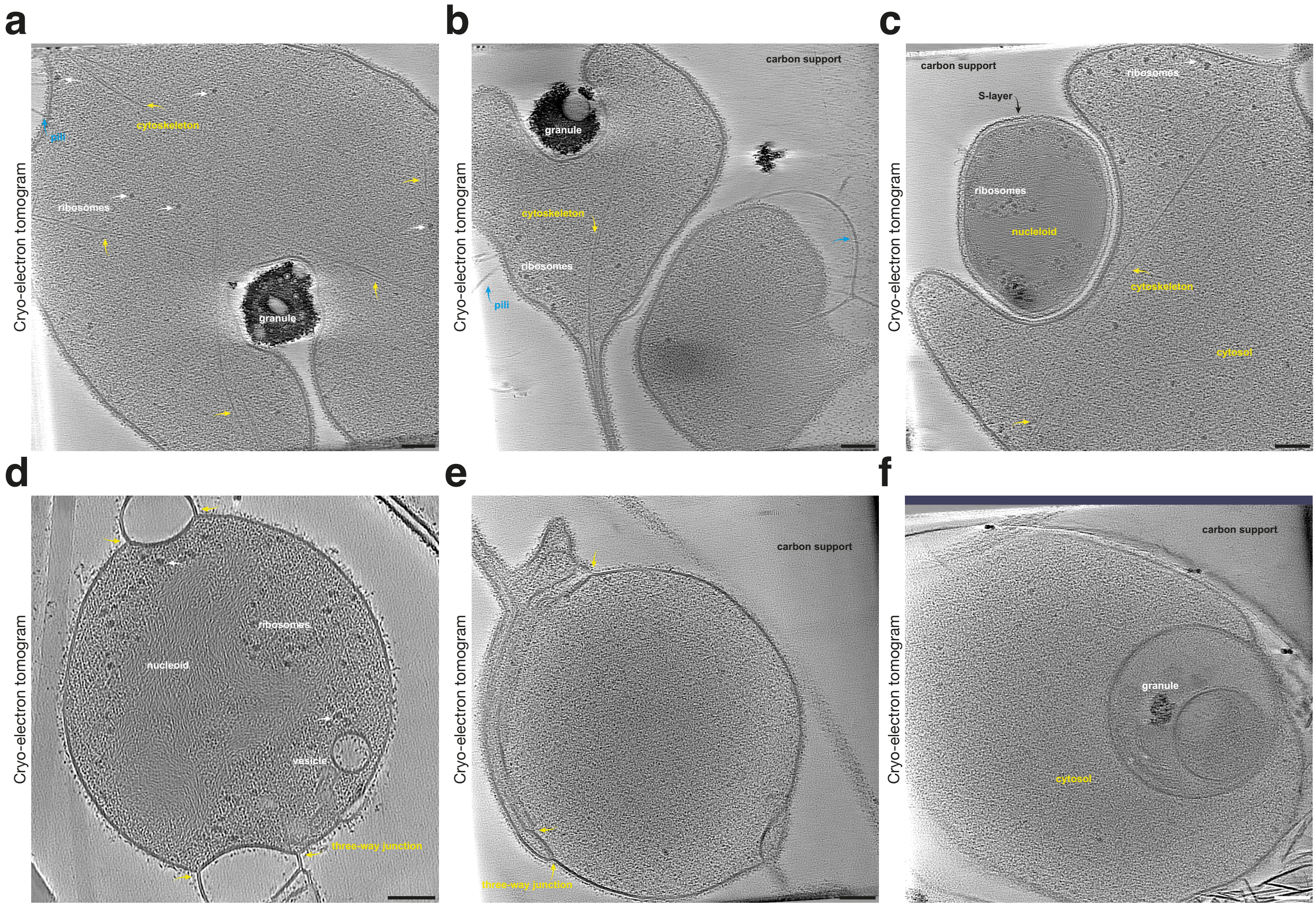


**Supplementary Figure 11: Cryo-ET of the more infrequent membrane deformations observed in *Ca.* Y. umbracryptum cells.**
**(a-b)** Cryo-ET slices through the cell body a) and protrusion b) of a granule situated in an invagination of the surface of an *Ca.* Y. umbracryptum cell (marked). **(c)** Cryo-ET slice through the cell body of *Ca.* Y. umbracryptum partially enwrapping another S-layer coated micro-organism. **(d-e)** Cryo-ET slices through cell bodies of *Ca.* Y. umbracryptum with three-way membrane junctions (yellow) in the plasma membrane. **(f)** Cryo-ET slices through a cell body of a rare late-log phase *Ca.* Y. umbracryptum cell with a large, internalized vesicle containing a dense granule (black). Scale bars: 100 nm.

**Supplementary Table 1:** Genome statistics

| Organism | Size (Mb) | Completeness (%) | Contamination (%) | 5S | 16S | 23S | tRNAs |
|---|---|---|---|---|---|---|---|
| *Ca*. Y. umbracryptum | 2.9 | 90.75 | 3.04 | - | + | + | 45 |
| *P. syntrophicum* | 4.3 | 92.65 (100) | 6.22 (0) | + | + | + | 46 |
| *Ca*. L. ossiferum | 6.0 | 91.26 (100) | 9.14 (0) | + | + | + | 44 |

**Supplementary Table 2:** Cryo-ET data collection, refinement, and validation statistics

| | Late-Log culture | | | | Early-Log culture | |
|---|---|---|---|---|---|---|
| **Data collection** | | | | | | |
| Microscope | | Titan Krios G4 | | Titan Krios G3 | Titan Krios G3 | |
| Detector | | Falcon 4i | | K3 (Gatan) | K3 (Gatan) | |
| Software | | SerialEM[106] | | SerialEM[106] | SerialEM[106] | |
| Magnification | | 42,000 | | 42,000 | 42,000 | |
| Voltage (kV) | | 300 | | 300 | 300 | |
| Slit width (eV) | | 10 | | 20 | 20 | |
| Defocus range (μm) | | -4 to -8 | | -5 to -8 | -5 to -8 | |
| Pixel size (Å) | | 3.01 | | 2.13 | 2.13 | |
| Total exposure ($e^-/Å^2$) | 140 | 140 | 140 | 120 | 122 | 120 |
| Exposure per tilt ($e^-/Å^2$) | ~3.4 | ~3.4 | ~3.4 | ~2 | ~2 | ~1.96 |
| Total number of tilts | | 41 | | 61 | 61 | |
| Frames per tilt-movie | | EER-format | | 10 | 10 | |
| Tilt increment | | 3° | | 2° | 2° | |
| Tilt-series scheme | | dose-symmetrical | | dose-symmetrical | dose-symmetrical | |
| Tilt range | | ±60° | | ±60° | ±60° | |
| Tilt-series collected | 31 | 56 | 58 | 21 | 28 | 25 |
| Tilt-series used | 27 | 48 | 51 | 9 | 24 | 18 |
| **Data processing** | | | | | | |
| Software tilt-series alignment | | ARETOMO2[109] | | ARETOMO2[109] | ARETOMO2[109] | |
| Software reconstruction | | RELION5.0[32] | | RELION5.0[32] | RELION5.0[32] | |
| Initial particle images (no.) | | 10,469 | | | 6,805 | |
| Final particle images (no.) | | 2,277 | | | 2,155 | |
| Final Box-size (px) | | 256 x 256 x 256 | | | 256 x 256 x 256 | |
| Pixel size final rec. (Å) | | 3.01 | | | 2.13 | |
| Map resolution (Å) | | 10.14 | | | 10.69 | |
| FSC threshold | | 0.143 | | | 0.143 | |
| Map resolution range (Å) | | 8.4-27.0 | | | 8.9-24.8 | |
| Map sharpening *B* factor ($Å^2$) | | none | | | none | |
| EMDB code | | XXXXX | | | XXXXX | |

**Supplementary Table 4:** Media Composition

| | | Hamelin Pool Media (HPM) | Hamelin Pool Salt (HPS) Buffer | Minimal Loki Media (MLM)[7] | Archaeal Salt Media (ASM) |
|---|---|---|---|---|---|
| Salts | | | | | |
| | NaCl | 50.7 g/L | 50.7 g/L | 20.7 g/L | 50.7 g/L |
| | $MgSO_4 \cdot 7H_2O$ | 13.3 g/L | 13.3 g/L | - | 13.3 g/L |
| | $MgCl_2 \cdot 6H_2O$ | 7.23 g/L | 7.23 g/L | 5.0 g/L | 7.23 g/L |
| | $CaCl_2 \cdot 2H_2O$ | 2.7 g/L | 2.7 g/L | 1.36 g/L | 2.7 g/L |
| | KCl | 1.4 g/L | 1.4 g/L | - | 1.4 g/L |
| | $NH_4Cl$ | - | - | 0.54 g/L | - |
| | $KH_2PO_4$ | 0.14 g/L | 0.14 g/L | 0.14 g/L | 0.14 g/L |
| | $Na_2CO_3$ | To pH 7.5 | To pH 7.5 | - | To pH 7.5 |
| | $NaHCO_3$ | - | - | To pH 7.5 | - |
| Growth substrate | | | | | |
| | Yeast extract | 0.05% | - | - | 0.05% |
| | Casein hydrolysate | 0.05% | - | 0.1% | 0.05% |
| Reducing agents | | | | | |
| | $Na_2S \cdot 9H_2O$ | 0.3 g/L | - | 0.3 g/L | 0.3 g/L |
| | L-cysteine HCl | 0.3 g/L | - | 0.3 g/L | 0.3 g/L |
| Antibiotics | | | | | |
| | Ampicillin | 0.001 g/L | - | 0.002 g/L | 0.001 g/L |
| | Streptomycin | 0.001 g/L | - | 0.002 g/L | 0.001 g/L |
| | Kanamycin | 0.0005 g/L | - | 0.002 g/L | 0.0005 g/L |
| Trace solutions | | | | | |
| | Vitamin 10[60] | 3 mL/L | - | - | 3 mL/L |
| | SL10 *TE[59] | 1 mL/L | - | - | 1 mL/L |
| | Se/W | - | - | 1 mL/L | - |
| | Se/W/Mo | 1 mL/L | - | | 1 mL/L |
| | Acidic *TE[7] | - | - | 0.5 mL/L | - |
| | Alkaline *TE[7] | - | - | 0.5 mL/L | - |

*TE: Trace elements.

**Supplementary Video 1:** Slices through cryo-electron tomogram of late stage/stationary phase *Ca.* Y. umbracryptum cell shown in Figure 2i followed by overlay of sematic segmentation shown in Figure 2j.

**Supplementary Video 2:** Slices through cryo-electron tomogram of late stage/stationary phase *Ca.* Y. umbracryptum cell shown in Figure 2k followed by overlay of sematic segmentation shown in Figure 2l.

**Supplementary Video 3:** Slices through cryo-electron tomogram of early-log phase *Ca.* Y. umbracryptum cell shown in Figure 3d followed by overlay of sematic segmentation shown in Figure 3e.
